# Senescent host-derived extracellular vesicles potentiate antitumor immunity via bystander T cell activation

**DOI:** 10.64898/2025.12.23.696148

**Authors:** Chenlu Yao, Chenhui Weng, Heng Wang, Bingbing Wu, Rong Sun, Huaxing Dai, Fang Xu, Qingzhen Han, Dong Zheng, Yumin Wu, Chao Wang

## Abstract

Extracellular vesicles (EVs) are key mediators of intercellular communication, and those from the plasma of aged individuals are typically viewed as drivers of aging or even cancer. Here, we uncover a latent, potent antitumor capacity inherent to EVs derived from aged hosts. Specifically, we demonstrate that extracellular vesicles from the plasma of aged mice (A-EVs) exhibit robust tumor-suppressive activity in young tumor-bearing recipients. Mechanistically, A-EVs engage a dual mode of action: they directly induce senescence in cancer cells, enhancing their immunogenicity. More importantly, they concurrently activate non-tumor-specific bystander CD8⁺ T cells within the tumor microenvironment via cross-presentation of pathogen-derived antigens, thereby reshaping the immunosuppressive tumor microenvironment. This synergistic interplay potently reinvigorates antitumor immunity and sensitizes tumors to PD-L1 checkpoint blockade. Our findings establish A-EVs as a novel class of endogenous immunomodulators with broad therapeutic potential for cancer immunotherapy.

## Introduction

Cancer remains a formidable challenge to human health, with the efficacy of many therapeutic strategies often being limited by the immunosuppressive nature of the tumor microenvironment (TME)^3, 4^. While strategies like immune checkpoint blockade have shown remarkable success, they are ineffective in a substantial proportion of patients, often referred to as cold tumors, where T cell infiltration and activity are insufficient^5^. Furthermore, the prevalence of bystander T cells within the tumor microenvironment, which are incapable of recognizing tumor-specific antigens^6^, represents a critical challenge. Therefore, leveraging these cells to redirect anti-tumor immunity and fundamentally reshape the immunosuppressive TME are urgently needed.

The relationship between aging and cancer is complex and paradoxical^7, 8^. While advanced age is the single greatest risk factor for the development of cancer^9^, the aged organism also accumulates a unique set of systemic factors and cellular experiences whose functional impact on tumor growth is not fully understood. Extracellular vesicles (EVs), as key mediators of intercellular communication, are likely carriers of these age-related signals^10, 11^. EVs are nano-sized, membrane-bound vesicles that carry proteins, RNAs, and other molecular cargo derived from their parent cells^12, 13^. Recent studies have shown that EVs derived from senescent cells carry specific senescence-associated secretory phenotype (SASP) factors or senescence-associated microRNAs, amplifying the transmission of senescence^14–16^ or promoting cancer^2^. Conversely, EVs isolated from young donors have been confirmed to transfer rejuvenating RNAs and proteins, which can reverse cellular senescence and alleviate age-related dysfunction^17^.

In a surprising contrast to the general view of EVs derived from senescent cells as a risk factor for promoting aging and cancer^1, 2^, our study uncovers a latent, potent anti-tumor capacity of these EVs. In this study, we report the unexpected discovery that extracellular vesicles derived from the plasma of aged mice (A-EVs) significantly inhibit tumor growth when administered to young, tumor-bearing hosts. We demonstrate that A-EVs employ a synergistic, two-pronged mechanism to achieve this effect. First, we uncovered a distinct mechanism that A-EVs have the capacity to activate bystander antigen-experienced CD8⁺ T cells (Tby) within the TME through the presentation of non-tumor antigens. This effectively mobilizes a population of non-tumor-specific bystander CD8⁺ T cells to contribute to the TME modulation by releasing effector cytokines. Second, we found that A-EVs directly induce cancer cells’ senescence within the tumor. This pro-senescent effect is coupled with a marked enhancement of tumor immunogenicity. Our findings demonstrate that A-EVs suppress tumors by synergistically linking enhanced tumor immunogenicity with the activation of bystander T cell immunity. This dual mechanism potently boosts the overall anti-tumor immune response that sensitizes tumors to PD-L1 blockade therapy.

## Results

### Preparation and characterization of A-EVs

We first collected and characterized the A-EVs from the peripheral blood of aged mice (18 months old) according to the established protocol^17^. To serve as a control, EVs were also isolated from young mice (6-8 weeks old) of the same strain, designated Y-EVs (**Fig. 1a**). Transmission electron microscope (TEM) and dynamic light scattering (DLS) analyses both revealed that A-EVs and Y-EVs exhibit a spherical morphology with a diameter of approximately 100 nm (**Fig. 1b–d**), confirming the successful isolation of a population consistent with small extracellular vesicles (sEVs). Following storage at 4°C, −20°C, −80°C, or a lyophilization-rehydration cycle, the collected A-EVs underwent minimal alteration in size, morphology, and protein composition, demonstrating their great stability (**supplementary Fig. 1**). To molecularly characterize A-EVs, we performed western blot and proteomic analyses. As expected, these analyses revealed a signature of cellular senescence, with A-EVs exhibiting significant enrichment of the classic senescence markers p16 and p21, along with numerous SASP factors, compared to Y-EVs (**Fig. 1e-f and supplementary Fig. 2**). Notably, we detected a marked upregulation of MHC class I molecules and associated factors involved in the antigen processing and presentation within A-EVs (**Fig. 1e-f and supplementary Fig. 2**), suggesting an elevated antigen presentation capacity of A-EVs compared to Y-EVs. Gene Ontology (GO) and Gene Set Enrichment Analysis (GSEA) also indicated an enrichment of obsolete aging and antigen-presenting machinery in A-EVs, compared with Y-EVs (**Fig. 1g-h**). These results align with prior work that defined a definitive senescence-associated molecular signature in A-EVs^18^.

**Fig. 1.**
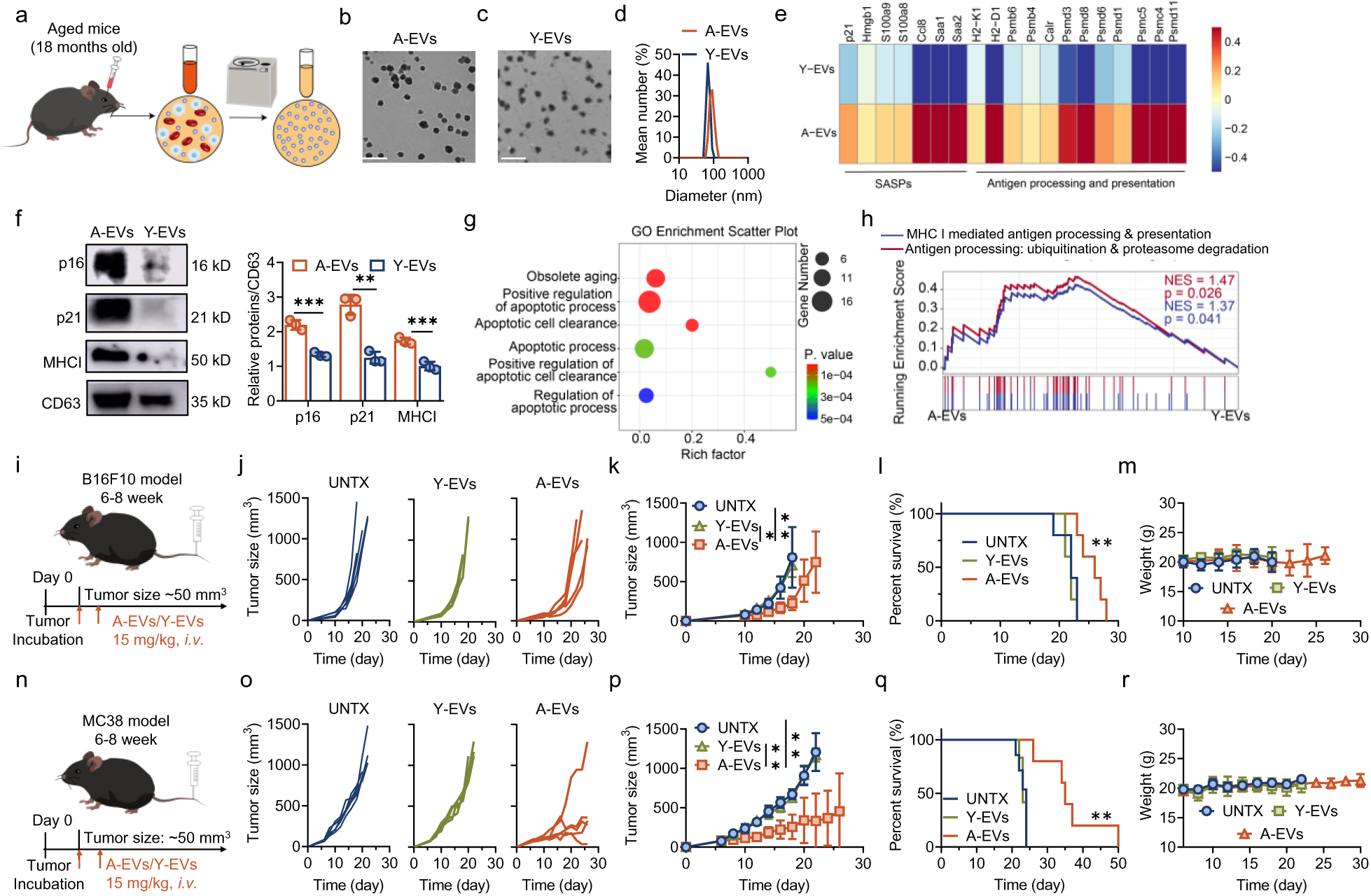
Preparation and characterization of A-EVs. **a**, Schematic diagram of the preparation of A-EVs from aged (18 months) mice. **b**, TEM image of A-EVs. Scale bar, 500 nm. **c**, TEM images of Y-EVs, scale bar, 500 nm. **d**, Dynamic light scattering of A-EVs and Y-EVs. **e**, Heatmap of proteomic analysis of A-EVs and Y-EVs. **f**, Representative results of western blot of A-EVs and Y-EVs, and corresponding quantitative analysis of p16, p21, MHC-I (n=3). **g**, Go enrichment pathway in proteomic analysis of A-EVs and Y-EVs. **h**, GSEA enrichment pathway in proteomic analysis of A-EVs and Y-EVs. **i**, Schematic diagram of intravenous treatment with A-EVs for B16F10 tumor-bearing mice. **j**, The tumor growth curves of B16F10 tumor-bearing mice in each group (n=5). **k**, Average curves of B16F10 tumor growth (n=5). **l**, Survival curves of B16F10 tumor-bearing mice (n=5). **m**, The weight curves of B16F10 tumor-bearing mice (n=5). **n**, Schematic diagram of intravenous treatment with A-EVs for MC38 tumor-bearing mice. **o**, The tumor growth curves of MC38 tumor-bearing mice in each group (n=5). **p**, Average curves of MC38 tumor growth (n=5). **q**, Survival curves of MC38 tumor-bearing mice (n=5). **r**, The weight curves of MC38 tumor-bearing mice (f n=5). Statistical significance was determined by Student’s t-test (two-tailed) and one-way ANOVA with the Tukey post hoc test. \*\*\**P* <0.001, \*\**P*<0.01. The data are presented as mean ±s.d.

### A-EVs delayed the tumor growth

Extracellular vesicles from aged individuals are known to accelerate aging primarily through the transfer of senescence-associated factors and the exacerbation of inflammatory responses^1^. However, the impact of A-EVs on tumor growth within tumor-bearing hosts remains unexplored. First, we found that A-EVs exhibited superior, integrin-dependent tumor accumulation after *i.v.* administration (**supplementary Fig. 3**). We next questioned the influence of A-EVs on tumor growth. Once subcutaneous B16F10 or MC38 tumors in young C57BL/6 mice (6-8 weeks old) reached approximately 50-60 mm³, the mice were treated with two intravenous injections of either A-EVs or Y-EVs (15 mg/kg). Strikingly, A-EVs from healthy aged donors functioned as potent antitumor agents in young tumor-bearing hosts, dramatically suppressing tumor growth and enhancing survival in both B16F10 and MC38 models. However, this therapeutic effect was absent in Y-EVs-treated mice (**Fig. 1i-r**) nor was it observed in A-EVs-treated aged hosts (**supplementary Fig. 4**). This robust efficacy, observed across distinct tumor types without significant side effects (**Fig. 1m, r**), demonstrates a broad-spectrum tumor therapeutic potential of A-EVs.

### A-EVs activate intratumoral effector CD8⁺ T cells directly

We next sought to determine the mechanism by which A-EVs inhibit tumor growth. Single-cell RNA sequencing (scRNA-seq) and flow cytometry were performed to characterize the tumor microenvironment post A-EVs treatment (**Fig. 2a-b**). The frequency of CD45^+^ immune cells was greater in the mice receiving A-EV than in the controls, as shown by scRNA-seq (**Fig. 2a**) and flow cytometry (**Fig. 2b**). To deeply profile the tumor immune microenvironment after A-EV treatment, we performed scRNA-seq on sorted CD45^+^ tumor-infiltrating cells. CD45^+^ cell population was divided into subsets according to the main markers of each immune cells (**supplementary Fig. 5a**), which included T cells (*Cd3e, Cd3d* and *Cd3g*), B cells (*Cd79a* and *Cd19*), DCs (*Itgax*), granulocytes (*Ly6g* and *Cxcr2*), macrophages (*Fcgr1* and *Adgre1*), monocytes (*Csf1r, Ly6c2* and *Ccr2*), and NK cells (*Klrb1c* and *Nkg7*). Among them, we discovered a marked increase in the frequency of T cells in the A-EV-treated tumor (**Fig. 2c**), which was further confirmed by flow cytometry (**Fig. 2d**). The frequency of B cells, granulocytes, DCs, monocytes, or macrophages did not change markedly. We then focused on the T cell subclusters to further divide them into several subsets according to each well-defined T cell marker, which included CD4^+^ T cells, CD8^+^ T cells, NK T cells, γδ T cells and other T cells (**Fig. 2e**, **and supplementary Fig. 5b**). The proportion and number of CD8^+^ T cells were the most prominently increased in the mice receiving A-EVs. We found a 2-fold increase in the fraction of CD8^+^ T cells within the CD3^+^ T cell population, which accounted for approximately 50% of the total T cells (**Fig. 2f and supplementary Fig. 6**). In addition, the expression of T cell dysfunction markers (*Entpd1*, *Lag3*, *Pdcd1*, *Ctla4* and *Tigit*) was reduced, whereas the expression of genes associated with cell proliferation (*Mki67*), memory (*Il2ra*), and cytotoxic effector molecules (*Gzmc*, *Gzmf* and *Ifng*) increased in CD8^+^ T cells post A-EV treatment compared to untreated mice (**Fig. 2g**). Gene set enrichment analysis (GO and KEGG) further corroborated these findings, showing significant enrichment for immune response, cell cycle, and MAPK and TNF signaling pathways in CD8^+^ T cells (**Fig. 2h-i**). Intriguingly, the transcriptional profile of A-EV-treated CD8^+^ T cells was significantly enriched in anti-viral and other pathogen-response KEGG pathways (*e.g.*, COVID-19, Hepatitis B, Measles) (**Fig. 2i**). This suggests that their potent activation state may also involve recognition of non-tumor antigens, such as those derived from endogenous viral elements or commensal pathogens, which could contribute to the overall immune activation. Crucially, tumor control was completely abrogated upon CD8^+^ T cell depletion, but not after depletion of CD4^+^ T cells or NK cells (**Fig. 2j-k and supplementary Fig. 7**), indicating that the antitumor effect of A-EVs may depend critically on CD8⁺ T cells.

**Fig. 2.**
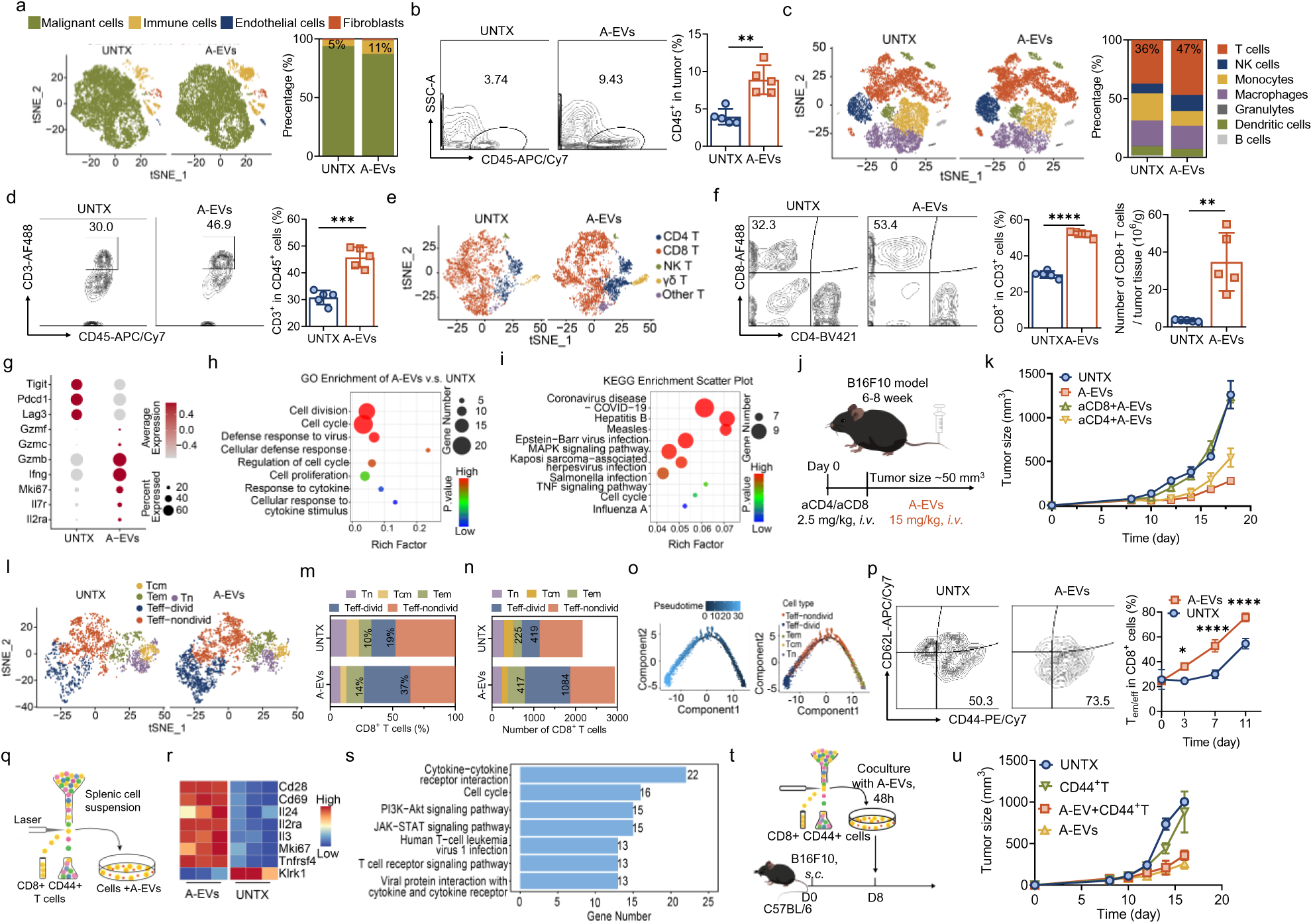
A-EVs activate intratumoral effector CD8^+^ T cells directly. **a**, T-SNE mapping of all cells in the tumor and relative proportions of cells in the tumor, including malignant cells, immune cells, endothelial cells, and fibroblasts. **b**, Representative flow cytometry plots of CD45^+^ immune cells in the tumor site after different treatments and the corresponding quantitative results (n=5). **c**, T-SNE mapping of all immune cells in the tumor sites after different treatments and relative proportions of immune cells in the tumor, including T cells, B cells, dendritic cells, granulocytes, macrophages, monocytes, and NK cells. **d**, Representative flow cytometry plots of CD3^+^ T cells in immune cells of the tumor site from different groups and the corresponding quantitative results (n=5). **e,** T-SNE mapping of T cells in tumors from different groups, including CD4^+^ T cells, CD8^+^ T cells, NK T cells, γδ T cells, and other T cells. **f**, Representative flow cytometry plots of CD8^+^ T cells in tumors of mice from the UNTX and A-EVs group and the corresponding quantitative results (n=5). **g**, Dot plot of genes’ differential expression in CD8^+^ T cells. **h**, GO enrichment scatter plot of CD8^+^ T cells in tumors. **i**, KEGG enrichment scatter plot of CD8^+^ T cells in tumors. **j**, Schematic diagram of antibody blockage for B16F10 tumor-bearing mice. **k**, Average curves of B16F10 tumor growth from mice receiving blockage therapy (n=3). **l**, T-SNE mapping of CD8^+^ T cells in tumors from mice in the UNTX and A-EVs group. **m**, Relative proportions of CD8^+^ T cells in the tumors, including naïve T cells (Tn), central memory T cells (Tcm), effector memory T cells (Tem), effector-dividing T cells (Teff-divid), and effector-nondividing T cells (Teff-nondivid). **n**, Absolute number of CD8^+^ T cells in the tumors. **o**, Pseudotime ordering of CD8^+^ T cells and naïve T cells, central memory T cells (Tcm), effector memory T cells (Tem), effector-dividing T cells (Teff-divid), and effector-nondividing T cells (Teff-nondivid). **p**, Representative flow cytometry plots of effector memory and effector T cells (Tem/eff) in tumors from mice in UNTX and A-EVs group and corresponding quantitative results of Tem/eff in CD8^+^ T cells over time (n=3). **q**, Schematic diagram of sorting CD8^+^ CD44^+^ T cells for *in vitro* activation experiment. **r**, Heatmaps of the expression of genes related to T cell activation from different *in vitro* treatment groups (n=3). **s**, KEGG enrichment bar plot of CD8^+^ CD44^+^ T cells from *in vitro* different treatments. **t**, Schematic diagram of the cell refusion experiment. **u**, Average curves of tumor growth of the cell refusion experiment (n=3). Statistical significance was determined by Student’s t-test (two-tailed) and one-way ANOVA with the Tukey post hoc test. \*\*\*\**P*<0.0001, \*\*\**P* <0.001, \*\**P*<0.01, \**P*<0.05, ns *P*>0.05. The data are presented as mean ±s.d.

To identify the most responsive CD8^+^ T cell subset, we further categorized them into five populations based on developmental origin and activation state: naïve T cells (Tn, *Tcf7, Lef1*), central memory T cells (Tcm, *Cd44, Sell*), effector memory T cells (Tem, *Cd44, Il7r*), nondividing effector T cells (Teff-nondivid, *Ifng, Tnf, Prf1, Mki67*^lo^) and dividing effector cells (Teff-divid, *Ifng, Tnf, Prf1, Mki67*^hi^) (**Fig. 2l-n and supplementary Fig. 8**)^19^. It is worth noting that the Tem and Teff-divid population demonstrated the most substantial expansion by A-EV treatment (**Fig. 2m-n**). Pseudotime analysis of the single-cell RNA-sequencing data also revealed that the proportions of Teff-divid cells increased along the trajectory in mice receiving A-EVs (**Fig. 2o**). Besides, results of flow cytometry also revealed the increase of effector memory/effector T cells (Tem/eff) (CD44^+^CD62L^-^) over time since the intravenous injection of A-EVs (**Fig. 2p**). This strongly indicated that A-EVs can induce the activation of CD44^+^ antigen (Ag)-experienced T cells^20, 21^ including both effector and memory T cells that have previously encountered antigen.

We next questioned whether A-EV activated these CD44^+^ CD8^+^ T cells directly. Splenic antigen-experienced CD8 T cells (CD8^+^CD44^+^) from healthy donor were sorted and co-cultured with A-EVs in the absence of tumor antigens (**Fig. 2q**). Intriguingly, exposure to A-EVs alone was sufficient to upregulate key genes involved in T cell activation and proliferation (e.g., *Cd28*, *Il2*, *Il2ra*, *Cd69*, *Mki67*) (**Fig. 2r and supplementary Fig. 9**). Consistently, KEGG pathway analysis demonstrated significant enrichment in cell cycle, PI3K-Akt signaling, JAK-STAT signaling, T cell receptor signaling pathway, and more importantly, some virus infection pathways (**Fig. 2s**). This demonstrates that A-EVs can directly activate memory CD8^+^ T cells, most likely through T cell receptor (TCR) recognition of antigens presented on A-EVs. These findings suggest that A-EV-reactive T cells represent a polyclonal memory population capable of recognizing a broad repertoire of non-tumor antigens, potentially enabling a generalized amplification of the anti-tumor immune response with a bystander effect. To prove the antitumoral efficacy of A-EV-primed CD44^+^ CD8^+^ T cells, we performed an adoptive transfer of these T cells into tumor-bearing mice (**Fig. 2t**). The mice that received A-EV–treated CD44^+^ CD8^+^ T cells exhibited a significant delay in tumor progression, demonstrating that the cells possessed enhanced antitumor function (**Fig. 2t-u and supplementary Fig. 10**). Collectively, these data indicate that A-EVs potently activate non-tumor antigen-experienced memory CD8^+^ T cells both transcriptionally and functionally, which contribute to the enhanced anti-tumor efficacy.

### Activation of bystander CD8^+^ T cells within the TME by A-EVs

Memory CD8^+^ T cells within the TME constitute a heterogeneous population, composed of both tumor antigen-experienced and non-tumor antigen-experienced (bystander) subsets^6^. The bystander subset originates from prior immune responses to unrelated pathogens or vaccinations and is subsequently recruited into the tumor stroma^22^. Notably, these bystander T cells frequently represent a substantial, and often dominant, proportion of the total tumor-infiltrating lymphocyte (TIL) pool. Memory CD8^+^ T cells are primarily activated by TCR signals, which are triggered by binding to epitope peptides presented by MHC-I. Given that A-EVs activated CD44^+^CD8^+^ T cells directly (**Fig. 2r-s**) and expressed high levels of MHC-I (**Figure 1e-h**), yet lacked tumor-associated antigens, we hypothesized that this response involved bystander activation of non-tumor-specific memory T cells. To provide more direct evidence, we employed an B16F10-OVA melanoma model, in which T cell specificity can be discerned via TCR recognition of the OVA antigen. Strikingly, TCR analysis of tumor-infiltrating CD8^+^ CD44^+^ T cells revealed that A-EV treatment preferentially expanded non-OVA-specific T cell clones, rather than OVA-specific clones (**Fig. 3a**). CD39 is a well-established marker to significantly enrich for tumor-specific T cell clones, particularly within the TME^23^. Based on the sequence of TCR and expression of *Entpd1* (the gene of CD39), we further classified CD8^+^CD44^+^ T cells into bystander (Tby), tumor-specific (Ttst), and other T cells (**Fig. 3b and supplementary Fig. 11-12**). Strikingly, A-EV treatment induced a pronounced expansion in the proportion and absolute number of Tby cells (**Fig. 3c**), particularly high-clonality Tby subsets (**Fig. 3d**), without significantly altering Ttst populations (**Fig. 3c–d and supplementary Fig. 13**). Notably, nearly all activated T cells belonged to the Tby compartment (**Fig. 3d**). These results clearly indicate that A-EVs directly promote bystander CD8⁺ T cell activation within tumors, dominating the T cell response without relying on tumor antigen specificity. Pathway analysis (KEGG and GO) of bystander T cells (Tby) within the CD8^+^ CD44^+^ T cells compartment also revealed their enrichment for defense responses to viral and other infectious diseases (**Fig. 3e-f**). This suggests that Tby cells likely represent a population of long-lived, pathogen-experienced memory T cells primed during earlier immune challenges. Functionally, these Tby cells in A-EV-treated tumors exhibited upregulated expression of key effector molecules, including the *Tnf*, *Il7*, *Gzmc,* and *Gzmf* (**Fig. 3g-h**), which was not observed in Ttst cells, reinforcing that A-EVs preferentially activate the Tby rather than the Ttst.

**Fig. 3.**
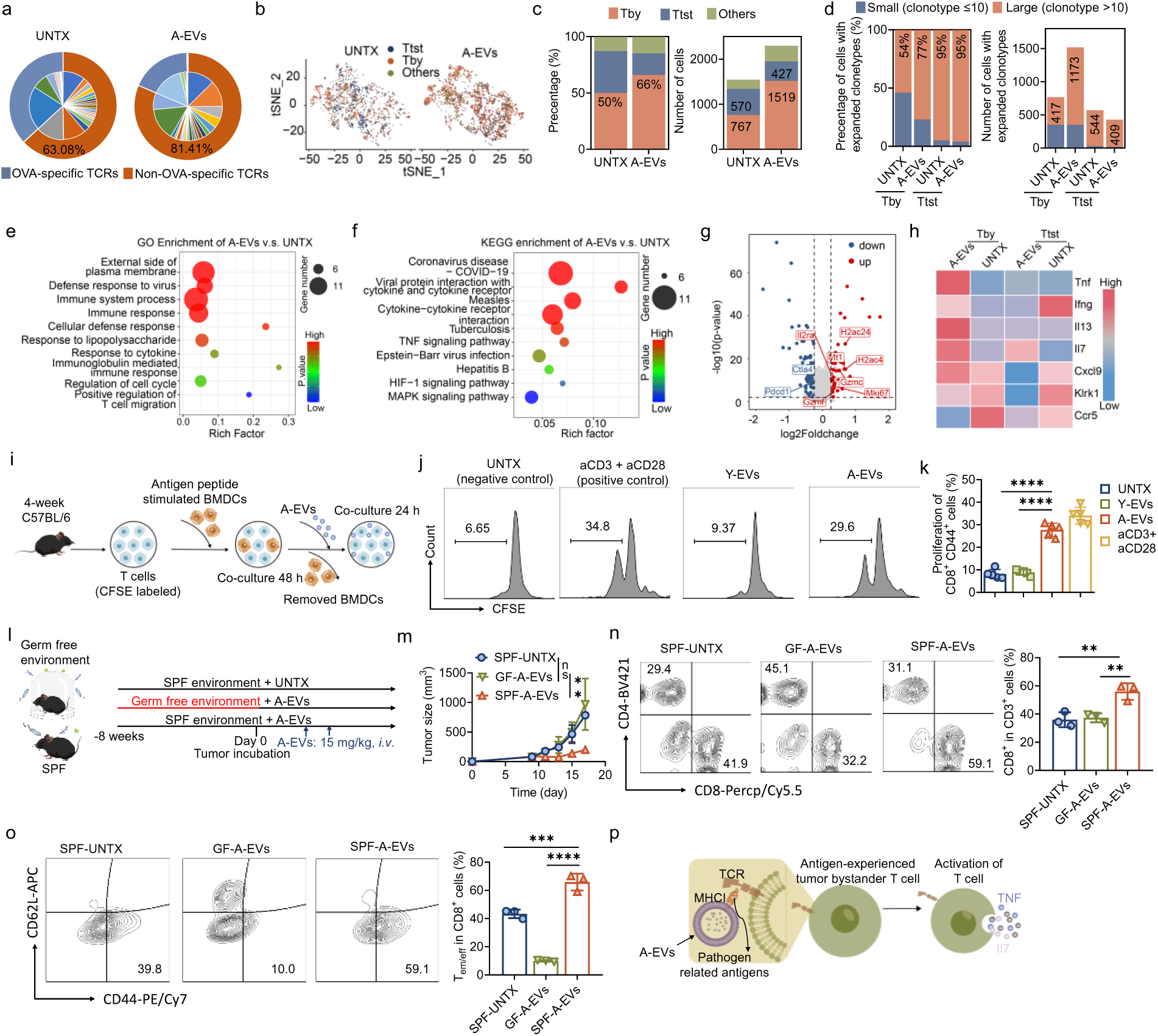
Activation of bystander T cells by A-EVs. **a**, Pie plot of TCR distribution of CD8^+^ CD44^+^ cells. **b**, T-SNE mapping of CD8^+^ CD44^+^ T cells. **c**, Relative proportions and absolute number of tumor-specific T cells (Ttst), bystander T cells (Tby), and other T cells in CD8^+^ CD44^+^ T cells. **d**, Frequencies of TCR clonotypes in Tby and Ttst CD8^+^ T cells from UNTX and A-EV-treated B16 tumors as indicated in **b**. **e**, GO enrichment scatter plot of Tby from CD8^+^ CD44^+^ T cells in tumors. **f**, KEGG enrichment scatter plot of Tby from CD8^+^ CD44^+^ T cells in tumors. **g**, Volcano plot of genes’ differential expression of Tby from CD8^+^ CD44^+^ T cells. **h**, Heatmap of genes related to cytokines in Tby and Ttst from CD8^+^ CD44^+^ T cells in tumors. **i**, Schematic diagram for the BMDC and T cell coculture model *in vitro*. **j**, Representative flow plots of CFSE^+^ in CD44^+^ T cells. **k**, The corresponding quantitative results of CFSE^+^ in CD8^+^ CD44^+^ T cells. **l**, Schematic diagram of the therapeutic design of mice living in different cleanliness levels. **m**, Average curves of B16F10 tumor growth (n=3). **n,** Representative flow plots of CD8^+^ T cells in tumors of mice from UNTX, GF-SPF, and SPF groups and the corresponding quantitative results (n=3). **o**, Representative flow cytometry plots of T_em/eff_ in tumors from mice in UNTX, GF-SPF, and SPF groups and corresponding quantitative results (n=3). **p**, Schematic diagram of bystander activation by A-EVs. Statistical significance was determined by Student’s t-test (two-tailed) and one-way ANOVA with the Tukey post hoc test. \*\*\*\**P*<0.0001, \*\*\**P* <0.001, \*\**P*<0.01, \**P*<0.05, ns *P*>0.05. The data are presented as mean ±s.d.

Additionally, memory CD8^+^ T cell activation can occur via either TCR engagement or cytokine-driven pathways, the latter characterized by NKG2D and CCR5 upregulation^24^. We found that A-EV treatment did not elevate these markers on Tby cells (**Fig. 3h**). This result further indicates that the observed Tby cell activation was mainly mediated by conventional TCR recognition of MHC-I presented on A-EVs, rather than by nonspecific cytokine stimulation.

The organism will be challenged by various pathogens during its growth. Based on our previous results and a recent study showing that senescent cells express viral antigens to enhance immune responses^25^, we hypothesized that A-EVs might present viral or bacterial antigens to activate bystander memory CD8^+^ T cells within the tumor microenvironment. To investigate this, T cells were first primed with BMDCs pulsed with antigens from influenza and tuberculosis — two common pathogens in murine infections. These pre-incubated T cells were then exposed to A-EVs (**Fig. 3i**). Intriguingly, we observed a significant expansion in CD8^+^ CD44^+^ T cells following co-culturing with A-EVs, but not Y-EVs (**Fig. 3j-k**). This finding directly demonstrates that A-EVs can present antigens specific to these common murine pathogens.

To further validate that bystander T cells activated by A-EVs were pathogen-experienced T cells, we utilized germ-free (GF) mice as a model (**Fig. 3l**). As GF mice are raised in a sterile environment with no pathogen exposure, they lack a pathogen antigen-experienced T cell pool, making them an ideal system to control for the effects of previous immune challenges. Strikingly, the antitumor efficacy of A-EVs was profoundly attenuated in GF mice compared to specific pathogen-free (SPF) live mice (**Fig. 3m and supplementary Fig. 14a-b**). This impaired efficacy was associated with a blunted T cell immune response in the tumor microenvironment, as evidenced by the unchanged proportions of CD3^+^ and CD8^+^ T cells between untreated and A-EV-treated GF mice (**Fig. 3n and supplementary Fig. 14c-e**). Furthermore, tumors from GF mice exhibited a near-complete absence of CD44^+^ CD8^+^ effector memory/effector T cells (Tem/eff) (**Fig. 3o**). All these results suggest that A-EVs could activate bystander antigen-experienced CD8^+^ Tby. This mechanism is dependent on the host’s pre-existing T cell pool (**Fig. 3p**), as the anti-tumor efficacy of A-EVs was profoundly attenuated in germ-free mice lacking such pathogen-experienced T cells.

### A-EVs induce tumor cellular senescence with a marked enhancement of tumor immunogenicity

In addition to bystander antitumor effect, previous studies have shown that senescent EVs act as “messengers”, thereby transmitting senescence signals to neighboring or distant cells and inducing a senescent phenotype^26^. We wonder whether the accumulation of A-EVs in tumor sites might directly induce senescence in tumor cells. To test this, we assessed changes in malignant cells within the tumor microenvironment following A-EV treatment. As expected, KEGG pathway analysis of malignant cells revealed marked upregulation of pathways associated with cellular senescence (**Fig. 4a and supplementary Fig. 15**). Conversely, the downregulation of key metabolic pathways—including oxidative phosphorylation and carbon metabolism—in A-EV-treated tumor cells (**Fig. 4a**) indicates that the treatment inhibited cancer cell proliferation. Furthermore, expression of senescence marker genes such as *Cdkn1a*, *Cdkn2a*, and *Trp53* was significantly elevated in tumor cells exposed to A-EVs (**Fig. 4b-d**), while the expression of these genes has no remarkable change in non-tumor cells within the TME **(supplementary Fig. 16**). Direct assessment of tumor tissue via β-galactosidase (β-Gal) staining showed a pronounced increase in senescent cells after A-EV treatment (**Fig. 4e**), and the upregulation of key senescence markers p16 and p21 was further confirmed at the protein level by immunofluorescence and western blotting (**Fig. 4f-h and supplementary Fig. 17**) compared to Y-EV-treated tumors, providing direct evidence that A-EVs induce senescence in the tumor microenvironment.

**Fig. 4.**
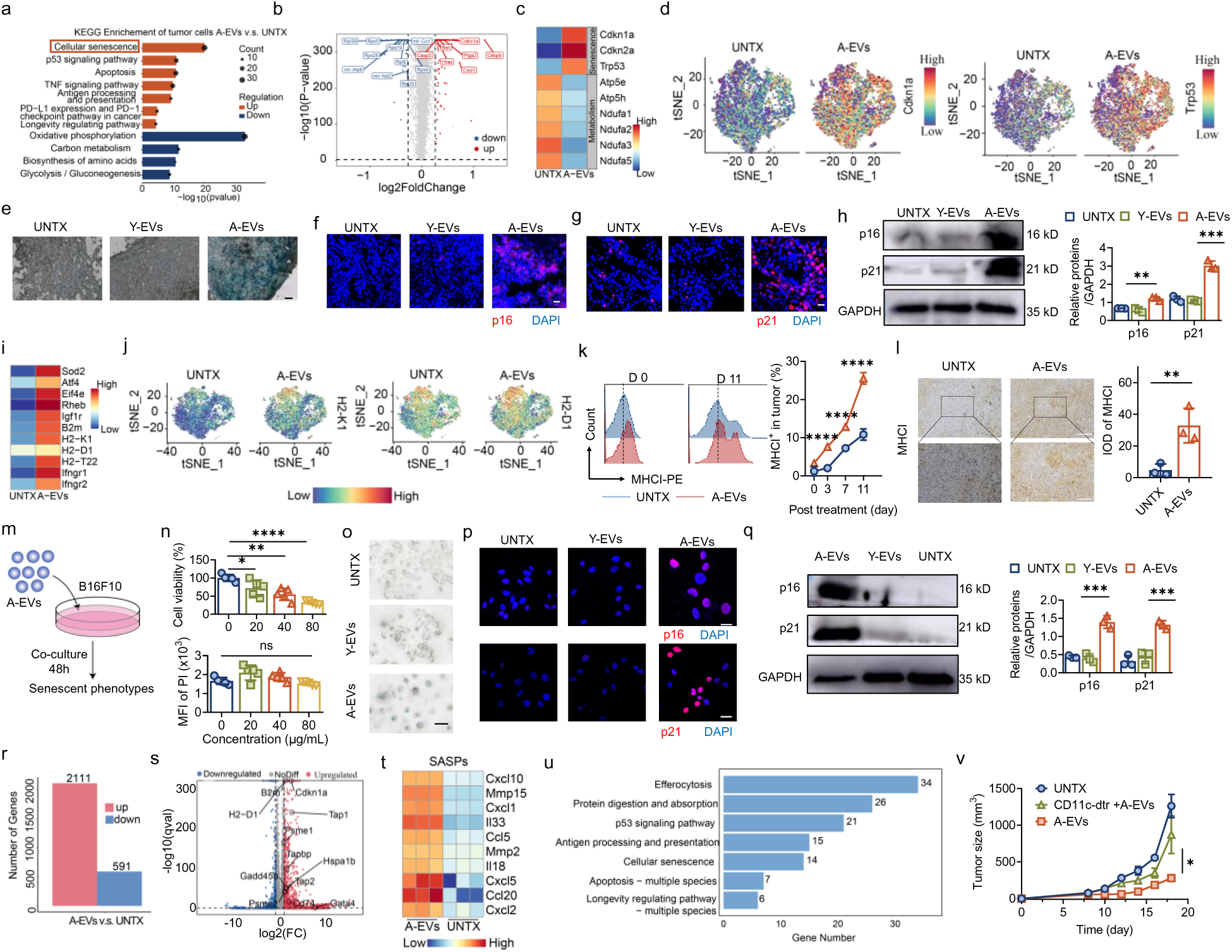
A-EVs induced the senescence of tumor cells both *in vivo* and *in vitro*. **a**, KEGG enrichment scatter plot of malignant cells. **b**, Volcano plot of genes differential expression in malignant cells. **c**, Heatmap of genes related to senescence and metabolism of malignant cells. **d**, Highlighted t-SNE plot of representative genes of *Cdkn1a* and *Trp53*. **e**, Representative images of SA-β-gal staining of tumor sections. Scale bar, 100 μm. **f**, Representative images of immunofluorescence staining of p16 (red) and DAPI (blue) in the tumor sections. Scale bar, 20 μm. **g**, Representative images of immunofluorescence staining of p21 (red) and DAPI (blue) in the tumor sections. Scale bar, 20 μm. **h,** Western blot of p16, p21, and GAPDH of tumors in different groups. Representative images and quantification results (n=3) are shown. **i**, Heatmap of genes related to antigen processing and presentation of malignant cells. **j**, Highlighted t-SNE plot of representative genes of *H2-K1* and *H2-D1*. **k**, Flow cytometry results of MHC-I expression on tumor cells in different groups. Representative images and quantitative results (n=3) are shown. **l**, Representative images of immunohistochemical staining of MHCI of tumor sections with different treatments (Scale bar: 100 μm, up; 50 μm, down) and quantitative results of immunohistochemical staining of MHC-I of tumor sections with different treatments (n=3). **m**, Schematic diagram of A-EVs cocultured with tumor cells *in vitro*. **n**, Cell viability was evaluated by MTT and quantitative results of PI staining of B16F10 cells with different dosages of A-EVs (n=5). **o**, Representative images of SA-β-gal staining of tumor cells in different groups. (Scale bar, 100 μm). **p**, Representative images of immunofluorescence staining of p16 and p21 (red), and DAPI (blue) of tumor cells in different groups. **q**, Western blot of p16, p21, and GAPDH of tumor cells in different groups. Representative images and quantification results (n=3) are shown. **r**, Bar plot of differential expression genes of tumor cells with different treatments. **s**, Volcano plot of differential expression of genes in tumor cells incubated with A-EVs in vitro and untreated tumor cells. **t**, Heatmap of genes related to SASPs of tumor cells. **u**, KEGG enrichment plot of tumor cells. **v**, Average curves of tumor growth in DC-depletion and wild-type B16F10 tumor-bearing mice (n=3). Statistical significance was determined by Student’s t-test (two-tailed) and one-way ANOVA with the Tukey post hoc test. \*\*\*\**P*<0.0001, \*\*\**P* <0.001, \*\**P*<0.01. The data are presented as mean ±s.d.

Unlike malignant cells, which often downregulate MHC-I to evade immunity, senescent cells upregulate antigen-presenting machinery^27, 28^. We thus investigated whether A-EV-induced senescence could restore MHC-I in tumors. Indeed, A-EV treatment significantly boosted MHC-I expression at both the transcriptional and protein levels (**Fig. 3i-l and supplementary Fig. 18a-e**). The concomitant upregulation of the interferon-gamma receptor (*Ifngr*) also suggested enhanced IFNγ sensitivity (**supplementary Fig. 18f**). These findings demonstrate that A-EVs enhance tumor immunogenicity and IFNγ responsiveness, potentially facilitating T cell-mediated clearance. To determine if A-EVs act directly, we treated B16F10 cells with A-EVs *in vitro* (**Fig. 4m**). This treatment directly suppressed proliferation (**Fig. 4n**) and induced a senescent phenotype, evidenced by elevated β-Gal activity (**Fig. 4o**) and increased p16/p21 expression (**Fig. 4p-q and supplementary Fig. 19**). RNA-seq and flow cytometry further confirmed the upregulation of senescence programs and antigen-presentation machinery (**Fig. 4r-u and supplementary Fig. 20**), demonstrating that A-EVs directly promote tumor cell senescence and enhance immunogenicity.

The induction of tumor cell senescence upon A-EV may increase tumor-specific T cells activation. Thus, A-EV, while effective in controlling tumor progression with bystander CD8 T cells alone, may also enhance the tumor-specific T cell response directly against tumors. Notably, tumor control with A-EV was better in wild-type compared with CD11c^+^ depleted mice (diphtheria toxin–treated CD11c-DTR transgenic mice), suggesting that Ttst also contributes to the antitumor efficacy of A-EV (**Figure 4v**).

### Aged human EVs inhibit tumor growth in mice

To test the translatability of our findings, we examined the effect of human aged EVs (hA-EVs) in a mouse tumor model. hA-EVs isolated from healthy human donors over 65 years old were administered to B16F10-tumor-bearing mice, with EVs from donors from 18-24 years old (hY-EVs) serving as controls (**Fig. 5a–g**). Similarly, hA-EVs significantly inhibited tumor growth, an effect not observed with hY-EVs (**Fig. 5h, i**). Mice treated with hA-EVs showed a marked improvement in survival, and the treatment was well-tolerated as evidenced by stable body weight (**Fig. 5j and supplementary Fig. 21a**). Consistent with murine findings, hA-EVs induced tumor senescence (β-gal, p16, p21) (**Fig. 5k–m**) and enhanced immunogenicity, evidenced by elevated MHC-I (**Fig. 5n**) and an activated bystander T cell phenotype—marked by expanded memory-derived and CD39^-^ CD8^+^T cells with reduced exhaustion (**Fig. 5o–q and supplementary Fig. 21b, c**). These results establish that the capacity of aging EVs to activate bystander T cell immunity and boost tumor immunogenicity is a conserved mechanism across species.

**Fig. 5.**
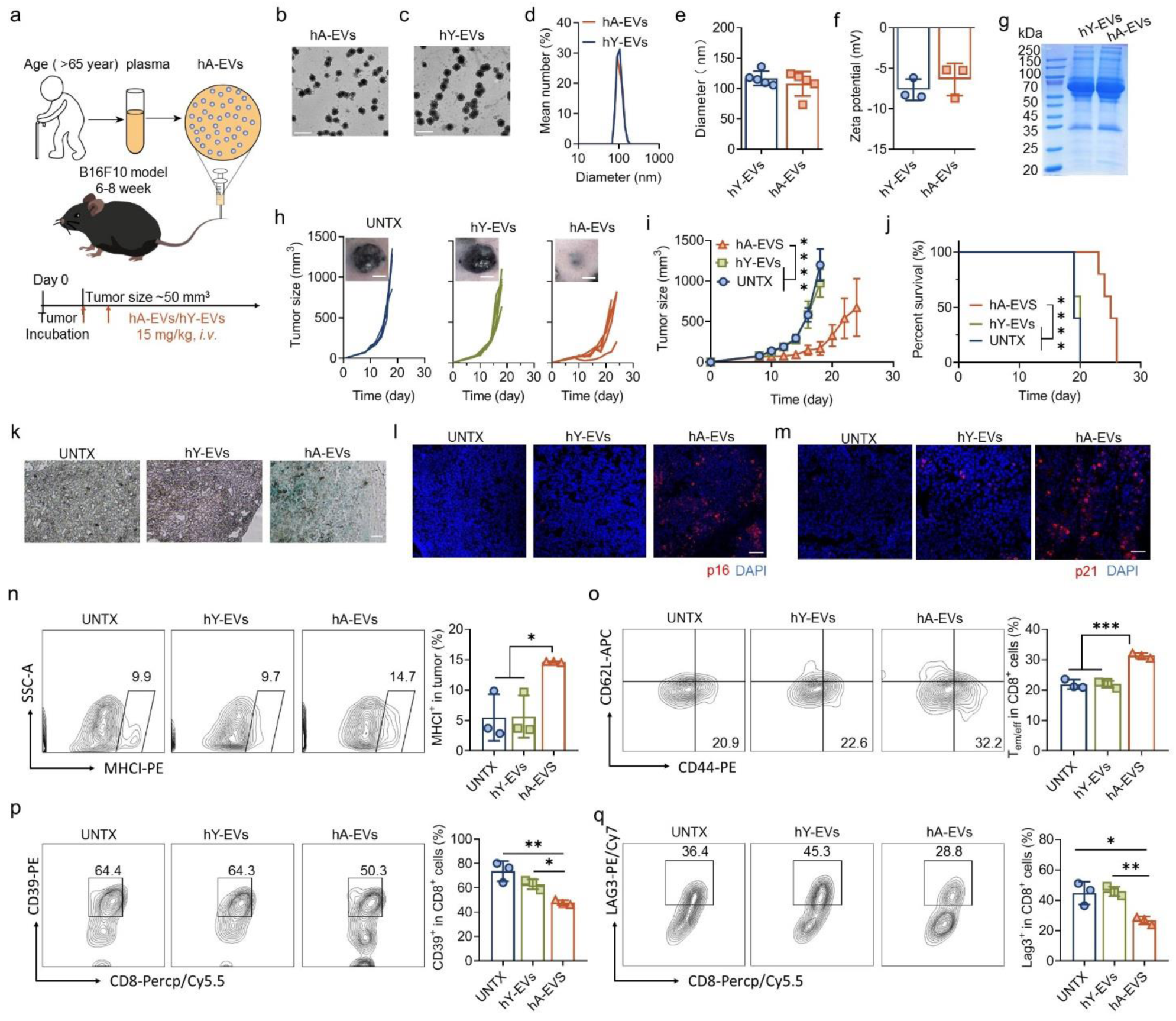
Human aged EVs inhibited tumor growth in mice. **a**, Schematic diagram of the preparation of hA-EVs and intravenous treatment of hA-EVs for B16F10 tumor-bearing mice. **b**, TEM image of hA-EVs. Scale bar: 500 nm. **c**, TEM image of hY-EVs. Scale bar: 500 nm. **d**, Dynamic light scattering of hA-EVs and hY-EVs. **e**, Diameters of A-EVs and Y-EVs detected by DLS (n=5). **f**, Zeta potential of hA-EVs and hY-EVs (n=3). **g**, SDS-PAGE of hA-EVs and hY-EVs. **h**, Representative images of tumors in B16F10-tumor-bearing mice (scale bar: 5 mm) and the tumor growth curves of B16F10 tumor-bearing mice in each group. **i**, Average curves of B16F10 tumor growth (n=5). **j**, Survival curves of B16F10 tumor-bearing mice (n=5). **k**, Representative images of SA-β-gal staining of tumor sections. Scale bar, 100 μm. **l**, Representative images of immunofluorescence staining of p16 (red), and DAPI (blue) in the tumor sections. Scale bar, 50 μm. **m**, Representative images of immunofluorescence staining of p21 (red), and DAPI (blue) in the tumor sections. Scale bar, 50 μm. **n**, Flow cytometry plots of MHCI in tumors of mice from different treatments and percentage of MHCI-positive tumor cells (n=3). **o**, Flow cytometry plots of effector memory and effector (Tem/eff) T cells among CD8^+^ T cells and percentage of effector memory and effector (Tem/eff) T cells among CD8^+^ T cells in tumor (n=3). **p**, Flow cytometry plots of CD39^+^ T cells among CD8^+^ T cells and percentage of CD39^+^ T cells among CD8^+^ T cells in tumor (n=3). **q**, Flow cytometry plots of Lag3^+^ T cells among CD8^+^ T cells and percentage of Lag3^+^ T cells among CD8^+^ T cells in tumor (n=3). Statistical significance was determined by one-way ANOVA with the Tukey post hoc test. \*\*\*\**P*<0.0001, \*\*\**P* <0.001, \*\**P*<0.01. The data are presented as mean ±s.d.

### A-EVs improve the antitumor efficacy of anti-PD-L1

Senescent tumor cells were previously reported to upregulate the expression of PD-L1 to escape immune evasion^29, 30^. Here, we also found an upregulation of PD-L1 in A-EVs-treated tumors over time (**Fig. 6a-c and supplementary Fig. 22a-b**). We therefore hypothesized that A-EVs, by enhancing tumor immunogenicity and activating bystander T cells, could effectively convert immunologically "cold" tumors into "hot" ones, thereby sensitizing them to PD-L1 checkpoint blockade. To test this, we combined A-EVs with an anti-PD-L1 antibody (aPD-L1) (**Fig. 6d**). In the poorly immunogenic B16F10 model, where aPD-L1 monotherapy was ineffective, the combination with A-EVs elicited marked tumor suppression and improved survival without overt toxicity (**Fig. 6e-h**). This enhanced efficacy was underpinned by a substantial increase in intratumoral T cell infiltration (**Fig. 6i-j and supplementary Fig. 22c-d**). The potent synergy was further validated in the MC38 mouse model, where the combination induced significant tumor regression and prolonged survival (**Fig. 6k-o**). Collectively, these results demonstrate that our strategy of using A-EVs to inflame the tumor microenvironment profoundly sensitizes tumors to PD-L1 blockade.

**Fig. 6.**
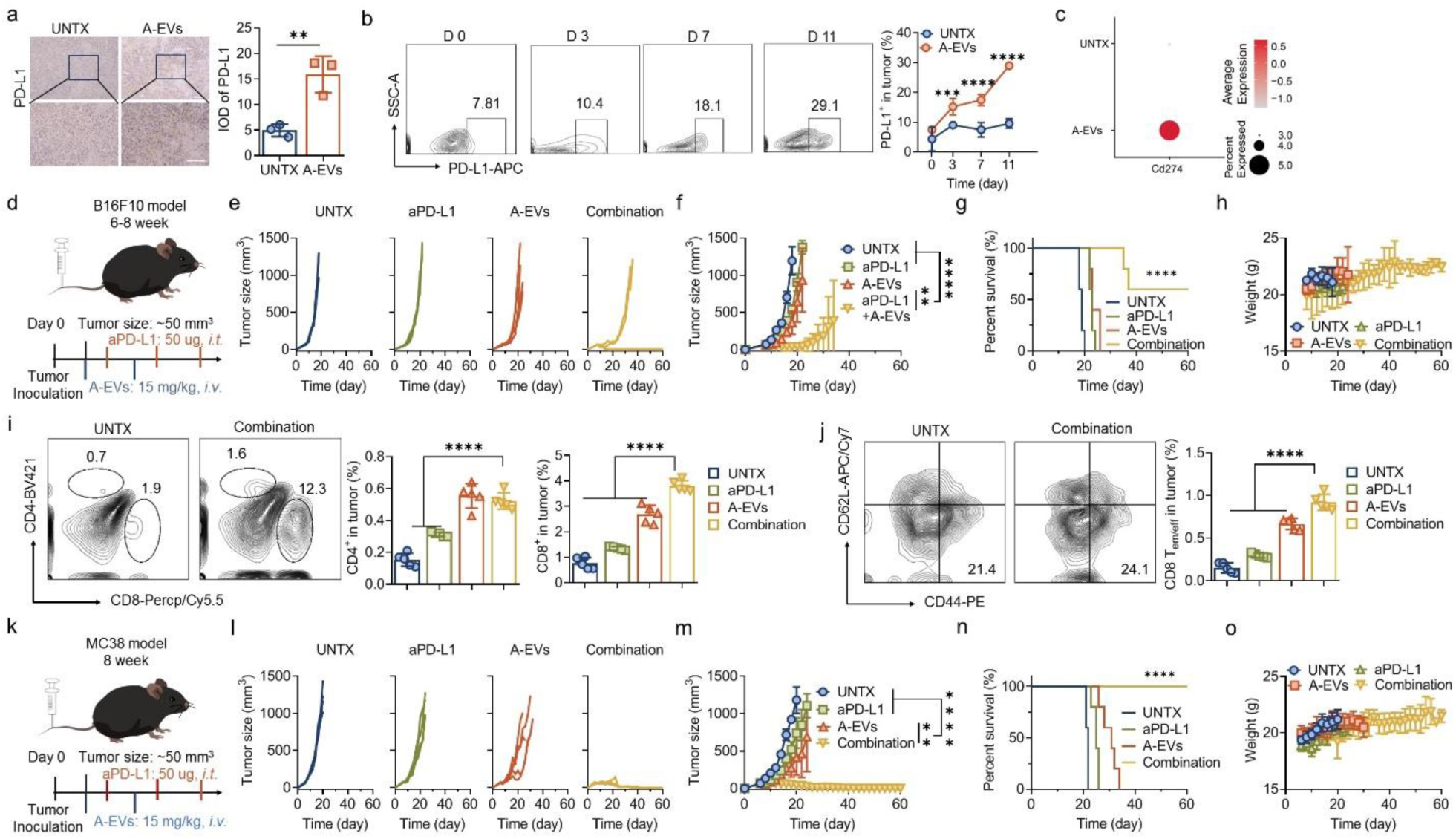
A-EVs enhanced the antitumor efficacy of anti-PD-L1. **a**, Representative images of immunohistochemical staining of PD-L1 of tumor sections with different treatments (Scale bar: 50 μm) and corresponding quantitative results (three tumor samples in each group). **b**, Flow cytometry plots of PD-L1 in tumors of mice from A-EV group over time, and corresponding quantitative analysis of PD-L1 level in tumors over time (n=3). **c**, Dot plot of the expression of *Cd274* in tumors from different groups in scRNA-seq. **d**, Schematic diagram of combination treatment with A-EVs and aPD-L1 for B16F10 tumor-bearing mice. **e**, Curves of tumor growth in mice after various treatments, including UNTX, aPD-L1, A-EVs, and combination **f**, Average tumor growth curves for each group (n=5). **g**, Survival curves of B16F10-tumor-bearing mice after different treatments. **h**, Weight curves of B16F10 tumor-bearing mice after various treatments (n=5). **i**, Representative flow cytometry plots of CD4^+^ and CD8^+^ T cells in tumors of mice, and the corresponding quantitative analysis of CD4^+^ and CD8^+^ T cells in tumors from four groups (n=5). **j**, Flow cytometry plots of effector memory and effector (Tem/eff) among CD8^+^ T cells and percentage of effector memory and effector (Tem/eff) among CD8^+^ T cells in tumors from four groups (n=5). **k**, Schematic diagram of combination treatment with A-EVs and aPD-L1 for MC38 tumor-bearing mice. **l**, Curves of tumor growth in mice after various treatments, including UNTX, aPD-L1, A-EVs, and combination. **m**, Average tumor growth curves for each group (n=5). **n**, Survival curves of MC38-tumor-bearing mice after different treatments. **o**, Weight curves of MC38 tumor-bearing mice after various treatments (n=5). Statistical significance was determined by one-way ANOVA with the Tukey post hoc test. \*\*\*\**P*<0.0001, \*\*\**P* <0.001, \*\**P*<0.01. The data are presented as mean ±s.d.

## Discussion

This study makes the paradigm-shifting discovery that extracellular vesicles from aged mice possess a potent latent anti-tumor capacity. The enhanced antitumor immunity mediated by A-EVs can be attributed to their unique biomolecular composition, which drives two mechanisms: Firstly, A-EVs activate non-tumor-specific "bystander" memory CD8^+^ T cells within the tumor microenvironment by presenting non-tumor antigens, mobilizing a previously overlooked T cell population for anti-tumor immunity. In our study, the activated bystander CD8^+^ T cells, likely primed by prior environmental pathogen exposure, secreted granzymes and effector cytokines within the tumor. Secondly, A-EVs directly induce senescence in cancer cells, and this pro-senescent effect is coupled with a significant enhancement of the tumor’s immunogenicity.

While cancer risk generally increases with age, a notable decline in cancer incidence occurs in the very oldest age groups (above 85), potentially due to biological aging processes that suppress tumor growth^31^. Cellular senescence is a protective response to accumulated stress or damage and plays a complex dual role in tumor biology^32, 33^. Recent studies have shown that cellular senescence initially acts as a tumor-suppressive mechanism by imposing irreversible cell cycle arrest^34^. Furthermore, senescent cells can remodel the microenvironment via the SASP^34^ or suppress oncogenes such as *Kras*^7^. Our hypothesis that extracellular vesicles from aging individuals could harbor anti-tumor capacity represents a novel extension of this classic tumor-suppressive paradigm. We propose that the senescent program may not only act cell-autonomously but could also be leveraged therapeutically. We demonstrate that A-EVs, as key messengers of senescence, not only directly induce senescence in cancer cells but also activate bystander memory CD8^+^ T cells within the tumor microenvironment (TME), thereby achieving synergistic TME remodeling.

Specifically, we discovered that A-EVs activate bystander memory CD8^+^ T cells through the presentation of viral or bacterial antigens (such as influenza and tuberculosis) (**Figure 3i-k**). This finding aligns with emerging evidence indicating that senescent cells—such as fibroblasts—can process and present viral antigens to cytotoxic T cells^35^ ^36, 37^. The underlying mechanism may involve compromised intracellular defense in senescent cells, which facilitates the reactivation of latent exogenous viruses^35^. Additionally, impaired virus-induced IFN-α production and downregulation of longevity-associated genes have been reported in senescence, collectively enhancing cellular susceptibility to infections (e.g., influenza) and potentially promoting viral replication^38^. These alterations are likely to promote the accumulation of viral or bacterial antigens on the surface of senescent cells. It follows that extracellular vesicles derived from such cells may carry these enriched antigens, thereby potentially amplifying anti-tumor immunity through bystander T cell activation.

Furthermore, our work provides a theoretical and experimental foundation for leveraging bystander T cell activation in cancer therapy. This concept—where non-tumor antigens trigger potent antitumor responses—is embodied by the activation of bystander T cells. It is established that a substantial proportion of tumor-infiltrating T cells are bystanders, which, while not recognizing tumor-specific antigens, can mount effective antitumor responses when exposed to pathogen-derived antigens^6, 39^. This recognition has motivated therapeutic strategies aimed at activating such bystander T cells. For example, administration of bacteria lacking tumor antigens has been shown to activate bacteria-specific CD8^+^ T cells, which subsequently enhance tumor-specific immunity and suppress tumor growth—particularly when a pre-existing antibacterial memory is present^40^. Similarly, vaccines carrying non-tumor cell epitopes (such as SARS-CoV-2 mRNA vaccines) have also been demonstrated to amplify antitumor immunity^41^. In addition, COVID-19 vaccination establishes a pool of SARS-CoV-2-specific memory T cells. This pre-existing immunity can be leveraged by redirecting these cells to recognize and attack tumors^42^. This mechanism represents a complementary form of immune surveillance, offering a promising strategy especially in contexts where tumor-specific T cells are scarce or exhausted. In line with these approaches, our study suggests that A-EVs—potentially enriched with pathogen-related antigens—may act as an endogenous “bystander activator,” thereby contributing to the robust antitumor immunity observed in our model.

In summary, we report a paradigm-shifting discovery: extracellular vesicles from aged mice possess a potent, previously unrecognized anti-tumor capacity. We establish that A-EVs orchestrate tumor regression through an unprecedented two-pronged mechanism—simultaneously activating non-tumor-specific bystander CD8^+^ T cells via cross-reactive antigen presentation and directly inducing senescence in cancer cells. These processes act synergistically to remodel the immunosuppressive tumor microenvironment and sensitize to PD-L1 checkpoint blockade therapy. Our findings not only challenge the conventional view of senescence-associated EVs but also introduce A-EVs as a novel class of endogenous immunomodulators with transformative potential for cancer therapy.

## Limitations of the study

For limitations, however, this study does not fully elucidate the specific molecular cargos within A-EVs responsible for senescence induction or T cell activation. *In vivo* biodistribution and long-term safety profiles of A-EVs also require further investigation.

## Materials and methods

### Materials

The used antibodies and other materials in this article are displayed in **Supplementary Table 1**.

### Cell lines

B16F10, MC38, and B16F10-OVA were purchased from the Cell Bank of the Shanghai Institutes for Biological Sciences, Chinese Academy of Sciences. MC38 were cultured in RPMI 1640 medium supplemented with 10% FBS (OPCEL, BS-1105), 1% penicillin (Seven, SC118-01), and 1% streptomycin (Seven, SC118-01). B16F10 and B16F10-OVA were cultured in DMEM medium with high glucose supplemented with 10% FBS, 1% penicillin, and 1% streptomycin. BMDCs were extracted according to previous literature^43^. Briefly, 6-8-week-old female C57BL/6 mice were euthanized. The bone marrow cells were extracted and cultured in RPMI 1640 medium supplemented with GM-CSF (20 ng/mL). The maturation of BMDCs was detected on day 6 after the extraction of the bone marrow cells. Cells were tested every 3 months to exclude the presence of mycoplasma. All the cells were cultured at 37 °C in 5% CO_2_ incubator. Authentication of cells was not performed after receipt.

### Animals

C57BL/6 mice (female, 6-8 weeks old) and germ-free C57BL/6 mice (female, 6-8 weeks old) were purchased from Changzhou Cavins Laboratory Animal Company. Mice were kept in the SPF animal facility with a 12-h light/dark cycle, constant room temperature (21 ± 1 °C), and relative humidity (40%-70%). Food and water were free to access. C57BL/6 mice were kept in the SPF animal facility until they reached 18 months of age (aged mice). Animal welfare and experimental procedures were carried out strictly under the supervision of the ethics committee of Soochow University. (Approval number, SUDA20240508A04).

### Preparation and characterization of EVs

Human whole blood was extracted from both young healthy donors (aged from 18 to 24 years) and old healthy donors (older than 60 years) by aseptic venipuncture and collected into a vacuum tube (∼5 ml) containing EDTA as an anticoagulant. Mouse whole blood was collected from the sinus vein of aged (18 months) and young (6-8 weeks) C57BL/6 female mice and resuspended in PBS, containing EDTA as an anticoagulant. Whole blood was centrifuged at room temperature for 15 min at 1600*g* to obtain the plasma. Then, the obtained plasma was diluted with PBS, followed by ultracentrifugation to prepare EVs^17^. The plasma was first centrifuged at 4 °C for 5 min at 500g, followed by 25 min at 3,000*g*, then at 4 °C for 60 min at 12,000*g*, and finally at 4 °C for 70 min at 120,000*g.* EVs were suspended in PBS and stored at -80 °C immediately.

The concentration of EVs was analyzed by quantifying the total protein content utilizing a BCA kit (Beyotime). The hydrated size and zeta potential of the EVs were evaluated by dynamic light scattering (DLS, Nano ZS90). The morphology of A-EVs and Y-EVs was acquired via transmission electron microscopy (TEM, FEI, TF20). The proteins on both A-EVs and Y-EVs were detected by SDS‒PAGE and analysed semi-quantitatively by Western blotting. In detail, A-EVs and Y-EVs were first lysed in RIPA lysis buffer (SW104-01) supplemented with 1 × proteinase inhibitor. After the samples were lysed for 30 min on ice, the mixture was centrifuged at 12000 rpm for 20 min, after which the total protein concentration was evaluated via a BCA kit (Beyotime). Total protein was separated via 12.5 % SDS‒PAGE, followed by transfer to a 0.45 μm PVDF membrane (IPVH00010, Merck). After that, the sections were incubated with rabbit anti-mouse p16 (1:1000), anti-p21 (1:1000), anti-CD63 (1:1000), anti-MHC-I (1:1000), or goat anti-rabbit (1:5000) antibodies. The fluorescence signals of these samples were detected by a DNA electrophoresis gel imager (AI600UV). For proteomics, protein samples of A-EVs and Y-EVs were extracted and quantified, followed by tryptic digestion and desalting. Then, liquid chromatography-tandem mass spectrometry (LC-MS/MS) analysis was performed using an Orbitrap Astral mass spectrometer (Thermo Scientific) operated in narrow-window data-independent acquisition (nDIA) mode. Raw MS data were processed using a sequence database search for peptide and protein identification.

### Cytotoxicity assessment of A-EVs *in vitro*

The cytotoxicity of A-EVs on tumor cells (B16F10) was detected by the MTT assay. Briefly, 5,000 cells were cultured in each well of 96-well plates. 12 h after the incubation, the supernatant was removed, and a new supernatant supplemented with different concentrations (0- 80 µg/mL) of A-EVs was added, and cells were cultured for another 24 h. Then, the supernatant was removed, 5 mg/mL MTT solution was added, and cultured for 4 h. After that, the MTT solution was aspirated, and 100 µL DMSO was added to each well. The absorbance at 570 nm of each well was detected with a microplate reader, after 10 minutes of shaking in the dark.

The viability of tumor cells (B16F10) after A-EVs treatment was detected by Propidium Iodide (PI) staining, followed by flow cytometry. Briefly, after the co-incubation with different concentrations (0-80 µg/mL) of A-EVs for 24 h, tumor cells were collected by centrifugation (2,000 rpm, 3 min), and resuspended in PBS. 1µL PI solution (1 mg/mL) was added to the cell suspension, followed by gentle mixing. The cells were stained for 15 minutes in the dark, and the fluorescence of PI was detected by flow cytometry immediately.

### The activation assessment of A-EVs on T cells *in vitro*

To evaluate the activation of A-EVs on antigen-experienced T cells, antigen peptides from Influenza (sequence: ASNENMETM, KAVYNFATM, Sangon) and Mycobacterium tuberculosis (sequence: TYQRTRALV, Sangon) as previously liturature^44^ were purchased, and stimulated BMDCs at a concentration of 10 μg/mL, for 24 h. T cells from 4-week-old female C57BL/6 mice were sorted by flow cytometry (BD, Asia), labeled with CFSE, and cocultured with antigen-presenting BMDCs for 48 h. Then, BMDCs were removed by centrifugation, and CFSE-labeled T cells were cocultured with Y-EVs or A-EVs (20 μg/mL) for another 48 h, while aCD3 (1 μg/mL) and aCD28 (5 μg/mL) were added as a positive control. After the incubation, the proliferation of T cells was detected by flow cytometry (BD, Asia) (anti-CD3-AF488, anti-CD8-Percp/Cy5.5, anti-CD44-PE/Cy7).

### Senescent induction of B16F10 cells by A-EVs *in vitro*

The expressions of senescence-associated β-Galactosidase (SA-β-Gal), p16, and p21 were detected 48 h after B16F10 cells were co-incubated with A-EVs and Y-EVs (20 µg/mL). For the expression of SA-β-Gal, a SA-β-Gal stain kit (Solarbio, G1580) was utilized. The expressions of p16 and p21 were observed by both immunofluorescence and Western blot. B16F10 cells with different treatments were first washed with PBS and fixed with 4% paraformaldehyde for 15 min at 37 °C. After that, cells were incubated with rabbit anti-mouse p16 (1:200) or p21 (1:200) overnight at 4 °C, followed by incubation with goat anti-rabbit AF594 (1:1000) for 2 h at room temperature. After that cells were stained with DAPI for another 15 min at room temperature and were observed by a confocal microscope (ZEISS, LSM 800). For Western blot, cells with different treatments were first lysed for 30 min on ice, the mixture was centrifuged at 12000 rpm for 20 min, after which total protein concentration was evaluated via a BCA kit (Beyotime). Total protein was separated via 12.5 % SDS‒PAGE, followed by transfer to a 0.45 μm PVDF membrane. After that, the sections were incubated with rabbit anti-mouse p16 (1:1000), anti-p21 (1:1000), anti-GAPDH (1:1000), or goat anti-rabbit (1:5000) antibodies. The fluorescence signals of these samples were detected by a DNA electrophoresis gel imager (AI600UV).

### RNA-sequencing of B16F10 and T cells cocultured with A-EVs

For RNA-sequencing of B16F10 cells and CD44^+^ CD8^+^ T cells, 1×10^6^ cells were collected for each sample, with three repeated samples in each group. Total RNA was isolated using TRIzol reagent (Thermo Fisher, 15596018) according to the manufacturer’s protocol. Samples with a concentration >50 ng/μL, RIN >7.0, and total mass >1 μg were selected for downstream processing. Poly(A)+ mRNA was enriched using oligo(dT) magnetic beads (Thermo Fisher, 25-61005) and subsequently fragmented using the NEBNext Magnesium RNA Fragmentation Module (E6150S) at 94°C for 5-7 minutes. First-strand cDNA was synthesized with SuperScript II Reverse Transcriptase (Thermo Fisher, 1896649), followed by second-strand synthesis using E. coli DNA Polymerase I (NEB, m0209) and RNase H (NEB, m0297) with dUTP incorporation for strand marking. The resulting double-stranded DNA was end-repaired, A-tailed, and ligated to adapters. Size selection was performed using magnetic beads. A strand-specific library was constructed through UDG digestion (NEB, m0280) and PCR amplification. Finally, paired-end sequencing (PE150) was performed on an Illumina Novaseq 6000 platform (LC Bio).

### *In vivo* Biodistribution of A-EVs in tumor-bearing mice

First, tumor-bearing mouse models were established. B16F10 tumor cells (2×10^6^) were injected subcutaneously into the right flanks of C57BL/6 mice. To investigate the contribution of integrins in tumor targeting of A-EVs, an integrin inhibitor (Cilengitide, MCE) was intraperitoneally injected into tumor-bearing mice (1 mg/kg, three times). When the tumor size reached about 100 mm³, free Cy5.5, Y-EVs-Cy5.5, and A-EVs-Cy5.5 were intravenously injected. 24 h after the injection, mice were sacrificed, and the fluorescence intensity of main organs, including heart, liver, spleen, lung, kidney, and the tumor, was observed by a small animal live fluorescence imaging system (Lumina III). The signaling intensity of tumors was also observed via a confocal microscope. Briefly, the tumors were first embedded in OCT medium and cut into 10-μm-thick slices utilizing a freezing microtome (Leica, CM1860). Samples were then fixed with 4% paraformaldehyde for 15 min at 37 °C, followed by the staining of DAPI for another 30 min at 37 °C. The samples were finally observed by a confocal microscope (Zeiss, LSM800).

### Treatment of subcutaneous tumors by intravenous administration of A-EVs

On day 0, B16F10 (2×10^6^) and MC38 (2×10^6^) tumor cells were injected into the right flanks of 6-8-week-old C57BL/6 mice. For each tumor model, mice were divided into three groups: UNTX, Y-EVs, and A-EVs, with 5 mice in each group. Y-EVs and A-EVs (15 mg/kg) were injected via the tail vein when the tumor volume reached approximately 50-60 mm^3^ for two times. The tumor size and mouse weight were measured every other day until the tumor volume reached 1500 mm³. The tumor volume was calculated as length*width^2^/2. To further reveal the immune microenvironment and senescent induction of tumors after the treatment of A-EVs, mice were euthanized on day 7 after the first treatment of A-EVs, and tumors were analyzed by flow cytometry, immunofluorescence, immunohistochemistry, and western blot.

To analyze the senescent induction of tumors, tumors were first embedded in OCT medium and cut into 10-μm-thick slices utilizing a freezing microtome (Leica, CM1860). Samples were then fixed with 4% paraformaldehyde for 15 min at 37 °C, followed by covering with 3% BSA solution to avoid unspecific reactions with antibodies. Then, samples were covered with p16 rabbit recombinant antibody or p21 rabbit antibody, respectively, overnight at 4 °C. Samples were then covered with AF594-HRP goat-anti-rabbit antibody for another 2 h at room temperature, followed by the staining of DAPI for another 30 min at room temperature. The samples were finally observed by a confocal microscope. Some other sections were stained by the SA-β-gal kit as mentioned above. Besides, the expression of p16 and p21 was also detected by Western blot. Tumors were first crushed utilizing a homogenizer and centrifuged at 12,000 rpm for 20 min. The supernatant was collected, and the total protein concentration of the supernatant was detected using by BCA kit. Then the proteins of the supernatant were separated by 12.5% SDS-PAGE, followed by transfer to a 0.45 μm PVDF membrane. After that, the sections were incubated with rabbit anti-mouse p16 (1:1000), anti-p21 (1:1000), anti-GAPDH (1:1000), or goat anti-rabbit (1:5000) antibodies. The fluorescence signals of these samples were detected by a DNA electrophoresis gel imager (AI600UV). Moreover, the expressions of MHC-I and PD-L1 on tumor cells were analyzed by flow cytometry on day 0, 3, 7, and 11 after the first treatment of A-EVs.

To reveal the change of tumor immune microenvironment, single cell suspensions of the treated tumors were prepared, and the change of various immune cells in the tumor site was analyzed via flow cytometry. In detail, the tumor tissues were first cut into 1-2 mm³pieces, ground by a tissue homogenizer, and filtered through a nylon strainer with 300 mesh. The pellet cells were then centrifuged at 2000 rpm for 3 mins and suspended in red blood cell lysis buffer for 5 mins on ice to remove red blood cells. After two washes of PBS, the cells were resuspended in 100 μL PBS, with FcR blocking regent (Biolegend, 1:200) and incubated on ice for 15 mins. Then, the cells were centrifuged at 2000 rpm for 3 minutes and suspended in PBS containing 3% BSA, and stained with surface antibodies. These included CD45 (CD45-APC/Cy7) immune cells, CD3 T (CD3-AF488) cells, CD8 T (CD3-AF488, CD8-Percp/Cy5.5) cells, CD4 T (CD3-AF488, CD4-BV421) cells, and CD8 Tm/eff (CD3-AF488, CD8-Percp/Cy5.5, CD44-PE/Cy7, CD62L-APC/Cy7^-^) cells. The exhaustion markers were also detected in CD8^+^ cells by flowcytometry, including PD-1 (CD8-Percp/Cy5.5, PD-1-APC), Lag3 (Lag3-PE/Cy7, CD8-Percp/Cy5.5), CD39 (CD39-PE, CD8-Percp/Cy5.5).

To evaluate the therapeutic efficacy of A-EVs in aged hosts, B16F10 (2×10^6^) tumor cells were injected to the right flanks of aged (18 months old) female C57BL/6 mice. Tumor-bearing aged mice were randomly divided into three groups: UNTX, Y-EVs and A-EVs, with 3 mice in each group. When the tumor volume reached 50-60 mm^3^, A-EVs or Y-EVs were intravenously injected for two times every other day, with a dose of 15 mg/kg. The tumor size and mouse weight were measured every other day until the tumor volume exceeded 1500 mm³.

In the treatment of hEVs, B16F10 tumor cells were injected into the right flank of 6-8-week-old C57BL/6 mice. Tumor-bearing mice were randomly divided into three groups, named UNTX, hY-EVs, and hA-EVs, with 5 mice in each group. hA-EVs or hY-EVs were injected intravenously when the average tumor volume was about 50 mm^3^ for two times. The tumor size and mouse weight were measured every other day until the tumor volume reached 1500 mm³. Senescent induction and the change of tumor immune microenvironment were also tested in human-EV-treated tumors, as mentioned.

In the combination therapy, B16F10 tumor cells (2×10^6^) and MC38 (2×10^6^) were injected into the right flanks of 6-8-week-old C57BL/6 female mice on day 0. For each tumor model, mice were divided into three groups, named UNTX, aPD-L1, A-EVs, and A-EVs+aPD-L1, with 5 mice in each group. A-EVs (15 mg/kg) were intravenously injected when the tumor volume reached approximately 50 mm^3^ for two times. aPD-L1 (50 μg) was intratumorally administered every other day for three times. The tumor size and mouse weight were measured every other day to evaluate the therapeutic efficacy of combination therapy until the tumor volume reached 1500 mm³. The infiltration of T cells was evaluated on day 21 by flow cytometry, stained by CD3-AF488, CD4-BV421, CD8-Percp/Cy5.5, CD44-PE, and CD62L-APC.

### Single-cell RNA sequencing and TCR analysis of the tumor site

A-EV-treated B16F10-OVA tumor and untreated B16F10-OVA tumor were collected on day 11 after the first treatment and were prepared into single-cell suspensions. Cell suspensions were adjusted to a concentration of 700–1200 cells/μL for encapsulation. Single cells, barcoded gel beads, and master mix were co-encapsulated into Gel Beads-in-Emulsion (GEMs) using a 10x Chromium controller. Within GEMs, reverse transcription was performed to synthesize barcoded cDNA. After GEM breakage, cDNA was amplified via PCR. For mRNA library construction, amplified cDNA was fragmented to 200–300 bp, and Illumina adapters were ligated, followed by PCR amplification. For immune repertoire (TCR) analysis, target regions were enriched using nested PCR with constant region primers, followed by fragmentation, end repair, A-tailing, and adapter ligation to construct sequencing-ready libraries. Finally, all libraries were sequenced on an Illumina platform using paired-end sequencing. Bioinformatics analysis was performed utilizing the OmicStudio tool (LC-Bio Technology) at https://www.omicstudio.cn/tool.

### Mechanism of intravenous administration of A-EVs for the treatment of subcutaneous tumors

To investigate the contribution of specific immune cells to the antitumor efficacy of A-EVs, *in vivo* depletion using neutralizing antibodies was performed. On day 0, B16F10 tumor cells (2×10^6^) were injected on the right flank of 6-8-week-old C57BL/6 female mice. On day 0 and day 3, CD8, CD4, and NK1.1 antibodies were intravenously injected into the tumor-bearing mice to neutralize the corresponding antibodies, with a dose of 50 μg per time per mouse. For the DC depletion mouse model, CD11c-dtr mice were injected intraperitoneally with 100 ng DT for three times as reported before^45^. To confirm the depletion of these immune cells, the peripheral blood of mice treated with antibodies and untreated mice was collected, and the percentage of CD8^+^ T cells, CD4^+^ T cells, and NK cells was detected by flow cytometry. When the tumor volumes reached about 50 mm³, A-EVs (15 mg/kg) were injected through the tail vein for two times every other day. Tumor size and weight of tumor-bearing mice were monitored every other day until the tumor volume exceeded 1500 mm³. The tumor volume was calculated according to the formula: Tumor volume = length ×width ×width/2.

To further evaluate the contribution of antigen-experienced (CD44^+^) CD8^+^ T cells, an adoptive transfer experiment was performed. On day 0, B16F10 tumor cells (2×10^6^) were inoculated on the right flank of 6-8-week-old C57BL/6 female mice. Mice were divided into 3 groups: UNTX, CD8^+^ CD44^+^ T, and A-EVs, with three mice in each group. On day 6, antigen-experienced (CD44^+^) CD8^+^ T cells from young and healthy mice were sorted by flow cytometry (BD, Asia), followed by coculturing with A-EVs (20 μg/mL) *in vitro*. After coculturing for 48 h, antigen-experienced (CD44^+^) CD8^+^ T cells were intravenously injected into tumor-bearing mice, with a dose of 1×10^6^ cells per mouse on day 8. At the same time, A-EVs (15 mg/kg) were intravenously injected for two times every other day. Tumor size and weight of tumor-bearing mice were monitored every other day until the tumor volume reached 1500 mm³. The tumor volume was calculated as follows: Tumor volume = length ×width ×width /2.

To investigate the importance of prior microbial exposure on the antitumor efficacy of A-EVs, the responses of germ-free (GF) mice with those of conventional mice were compared. 8-week-old mice raised in a germ-free (GF) environment from birth were purchased and then raised in a SPF environment during the experiment. On day 0, B16F10 tumor cells (2×10^6^) were inoculated on the right flank of 8-week-old C57BL/6 female mice. When the tumor volumes reached about 50-60 mm³, A-EVs (15 mg/kg) were intravenously injected for two times every other day. Mice were divided into three groups: SPF-UNTX, SPF-A-EVs, and GF-A-EVs, with three mice in each group. Tumor size and weight of tumor-bearing mice were monitored every other day until the tumor volume reached 1500 mm³. The tumor volume was calculated as follows: Tumor volume = length ×width ×width /2. The tumor immune microenvironment was detected by flow cytometry, staining by CD3-AF488, CD4-BV421, CD8-Percp-Cy5.5, CD44-PE/Cy7, and CD62L-APC.

### Biosafety assessment of intravenous injection A-EVs

To analyze the biosafety of A-EVs in a mouse model, 6-8-week-old female C57BL/6 mice were injected with A-EVs or Y-EVs (15 mg/kg) intravenously for two times every other day. 10 days after the first injection of A-EVs or Y-EVs, mice were euthanized, and the main organs, including heart, liver, spleen, lung, and kidney, were collected for the detection of senescent induction. The expression of p21 in different organs was analyzed by immunofluorescence, and the expression of β-gal was analyzed by the SA-β-gal kit. Besides, the main organs were also collected for Hematoxylin and Eosin (H&E) Staining in the A-EV-treated group and the untreated group. Moreover, the blood biochemical analysis of mice injected with A-EVs and untreated mice was utilized to evaluate the injury of main organs, including alanine aminotransferase (ALT), aspartate aminotransferase (AST), UREA, creatinine (CREA), and creatine kinase (CK).

## Statistical analysis

All the results were statistically analyzed using GraphPad Prism 8 software and presented as the mean ±standard deviation (s.d.). Biological replicates were used in all experiments unless stated otherwise. Differences among more than two groups were evaluated using one-way ANOVA with Tukey’s multiple comparisons test. For comparisons between two groups, a two-tailed Student’s *t*-test was applied. In the analysis of diabetes incidence, log-rank tests were conducted to determine significant differences. Significant differences in all graphs were presented as \*\*\*\**P* < 0.0001; \*\*\**P* < 0.001; \*\**P* < 0.01; \**P* < 0.05; ns *P*>0.05. All the experiments were repeated at least three times.

## Materials availability

This study did not generate new unique reagents.

## Data and Materials Availability

The data needed to evaluate the findings of this study are available in the paper or in the supplementary material of this article.

## Acknowledgments

This work was supported by the National Natural Science Foundation of China (grant no. 32371476) and the Natural Science Foundation of the Higher Education Institutions of Jiangsu Province, China (grant no. 22KJA180003). This work was partly supported by the Collaborative Innovation Center of Suzhou Nano Science & Technology, the Priority Academic Program Development of Jiangsu Higher Education Institutions (PAPD), and the 111 Project.

## Author Contributions

C.W., Y. W., and C.Y. designed the project. C.Y. performed the experiments, collected the data, analyzed and interpreted the data, and wrote the first version of the paper. All authors contributed to the discussion of the results and implications, and the editing of the paper at all stages.

