## Supplementary information for "Senescent host-derived extracellular vesicles potentiate antitumor immunity via bystander T cell activation"

**Supplementary Table 1.** Materials applied in this study.

| Antibodies | Manufacturer | Catalog Number | Dilutions | Clone Numbers | Lot number |
|---|---|---|---|---|---|
| PE anti-mouse I-A/I-E | Biolegend | 107626 | 1:200 | M5/114.15.2 | B384156 |
| PE anti-mouse NK1.1 | Biolegend | 108708 | 1:200 | PK136 | B367623 |
| BV510 anti-mouse NK1.1 | Biolegend | 108738 | 1:200 | PK136 | B427713 |
| AF488 anti-mouse CD3 | Biolegend | 100210 | 1:200 | 17A2 | B433853 |
| BV421 anti-mouse CD4 | Biolegend | 100544 | 1:200 | RM4-5 | B446917 |
| Percp/Cy5.5 anti-mouse CD8a | Biolegend | 100734 | 1:200 | 53-6.7 | B434030 |
| APC anti-mouse CD62L | Biolegend | 104412 | 1:200 | MEL-14 | B433862 |
| PE anti-mouse CD44 | Biolegend | 103030 | 1:200 | IM7 | B438628 |
| APC/Cy7 anti-mouse CD62L | Biolegend | 104428 | 1:200 | MEL-14 | B417448 |
| PE/Cy7 anti-mouse CD44 | Biolegend | 103030 | 1:200 | IM7 | B438628 |
| APC/Cy7 anti-mouse CD45 | Biolegend | 147718 | 1:200 | I3/2.3 | B453145 |
| AF488 anti-mouse CD8a | Biolegend | 100723 | 1:200 | 53-6.7 | B400877 |
| APC anti-mouse PD-L1 | Biolegend | 124311 | 1:200 | 10F.9G2 | B418357 |
| APC anti-mouse CD8a | Biolegend | 100712 | 1:200 | 53-6.7 | B420169 |
| FITC anti-mouse CD3 | Biolegend | 100204 | 1:200 | 17A2 | B453826 |
| PE anti-mouse CD4 | Biolegend | 100408 | 1:200 | GK1.5 | B446624 |
| PE anti-mouse CD39 | Biolegend | 143803 | 1:200 | Duha59 | B402935 |
| PE anti-mouse PD-1 | Biolegend | 135205 | 1:200 | 29F.1A12 | B416225 |
| PE/Cy7 anti-mouse Lag3 | Biolegend | 125226 | 1:200 | C9B7W | B415271 |
| FITC anti-mouse CD45 | Biolegend | 157213 | 1:200 | S18009F | B400827 |
| InVivoMAb anti-PD-L1 | BioXcell | BE0101 | Diluted to 1 mg/mL | 10F.9G2 | 945124M2 |
| InVivoMAb anti-mouse CD8a | BioXcell | BE0004-1 | Diluted to 1 mg/mL | 53-6.7 | 79281J3 |
| InVivoMAb anti-mouse CD4 | BioXcell | BE0003-1 | Diluted to 1 mg/mL | GK1.5 | 728319M2 |
| InVivoMAb anti-mouse NK1.1 | BioXcell | BE0036 | Diluted to 1 mg/mL | PK136 | 828624M2 |
| CD9 Rabbit Recombinant Antibody | Proteintech | 84801-1 | 1:1000 | 242432B7 | 23014295 |
| p16 INK4a Rabbit Recombinant Antibody | ImmnuoWay | YM8152 | 1:1000 | PT0242R | 541684 |
| p21 Rabbit Recombinant Antibody | Proteintech | 28248-1 | 1:1000 | 3N14 | 00143521 |

| | | | | | |
|---|---|---|---|---|---|
| MHC Class I Rabbit Recombinant mAb | Cell Signaling Technology | 76828T | 1:1000 | E8E7N | 1 |
| Recombinant anti-GAPDH Antibody (HRP Conjugated) | Servicebio | ZB15004 | 1:1000 | | AC231104001 |
| HRP-Goat anti-rabbit Recombinant Secondary Antibody | Proteintech | SA00001 | 1:5000 | | 20001529 |

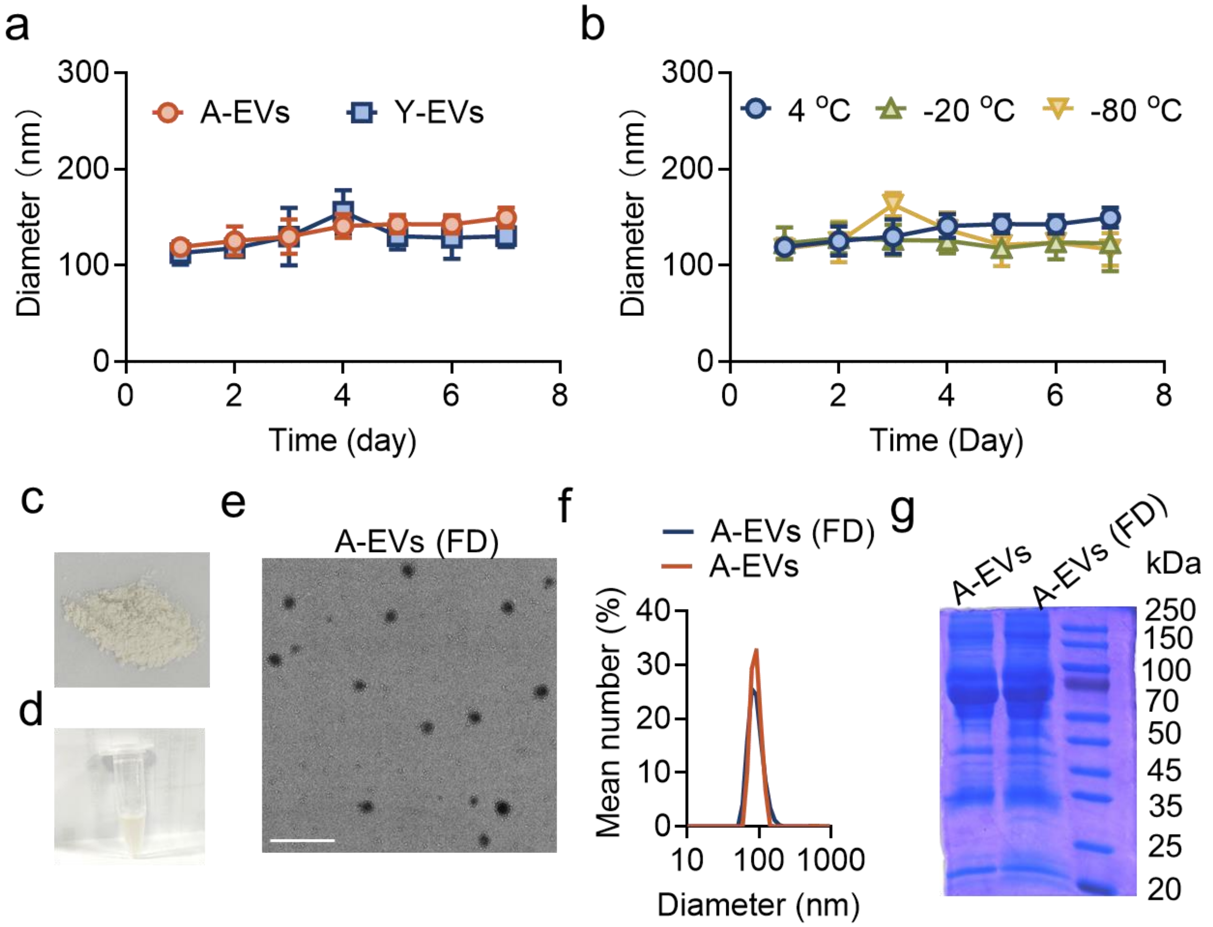


**Figure S1**. Characterization of A-EVs and Y-EVs. a, Diameter curves of A-EVs and Y-EVs within 7 days storage at 4 ºC (n=3). b, Diameter curves of A-EVs within 7 days storage at 4 ºC, -20 ºC, and -80 ºC (n=3). c, Photograph of A-EV lyophilized powder. d, Photograph of A-EV lyophilized powder redissolved in PBS. e, TEM image of A-EV Lyophilized powder redissolved in PBS. Scale bar, 500 nm. f, Dynamic light scattering of A-EVs and A-EV lyophilized powder redissolved in PBS. g, SDS-PAGE of A-EVs and A-EV lyophilized powder redissolved in PBS. The data are presented as mean ±s.d.

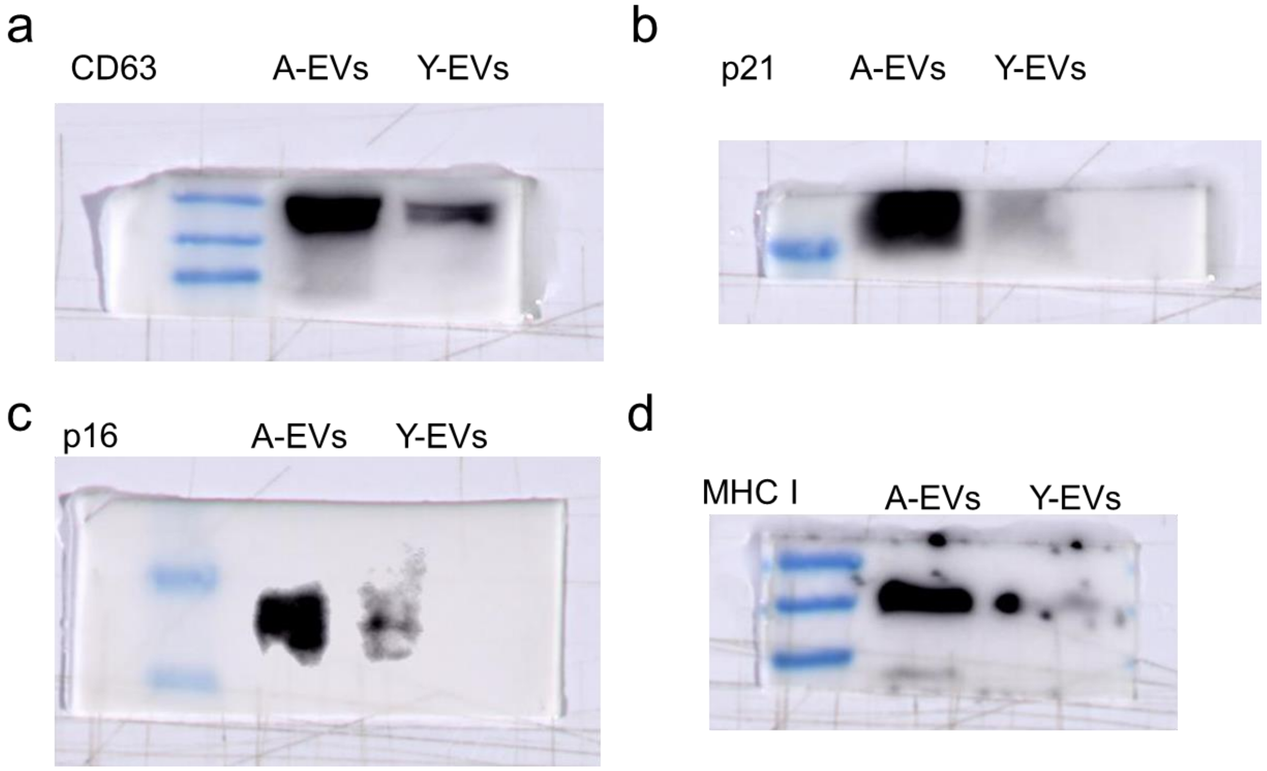


**Figure S2**. Western blot analysis of (a) CD63, (b) p21, (c) p16, and (d) MHC-I in A-EVs and Y-EVs.

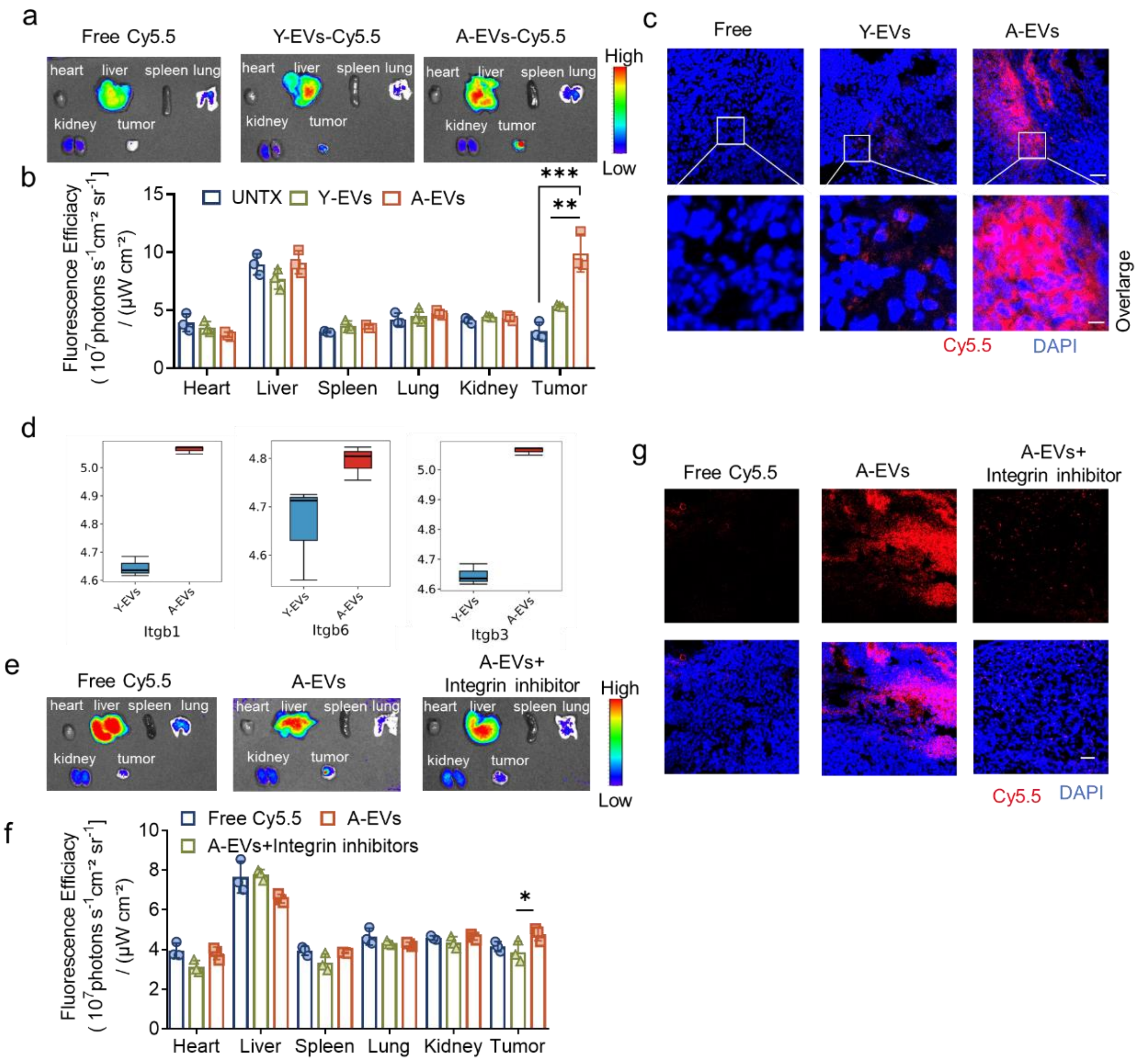


**Figure S3**. Biodistribution of A-EVs in B16F10 tumor-bearing mice. a, *Ex vivo* fluorescence images of major organs and tumors in B16F10 tumor-bearing mice with the intravenous injection of free Cy5.5, Y-EVs-Cy5.5, and A-EVs-Cy5.5 24 hours after the injection. b, Quantitative results of *ex vivo* fluorescence of main organs and tumor, shown in (a) (n=3). c, Confocal fluorescence images of B16F10 tumors after different treatments. Blue: DAPI, red: Cy5.5. Scale bar: 50 μm (up), 20 μm (down). d, Bar plots of the expression of integrins of A-EVs and Y-EVs. e, *Ex vivo* fluorescence images of major organs and tumors in B16F10 tumor-bearing mice with the intravenous injection of free Cy5.5, A-EVs, and A-EVs with integrin inhibitors, 24 hours after the injection. f, Quantitative results of *ex vivo* fluorescence of main organs and tumor, shown in (e) (n=3). g, Confocal fluorescence images of B16F10 tumors after different treatments. Blue: DAPI, red: Cy5.5. Scale bar: 50 μm. Statistical significance was determined by one-way ANOVA with the Tukey post hoc test. ***$P$ <0.001, **$P$<0.01, *$P$<0.05. The data are presented as mean ±s.d.

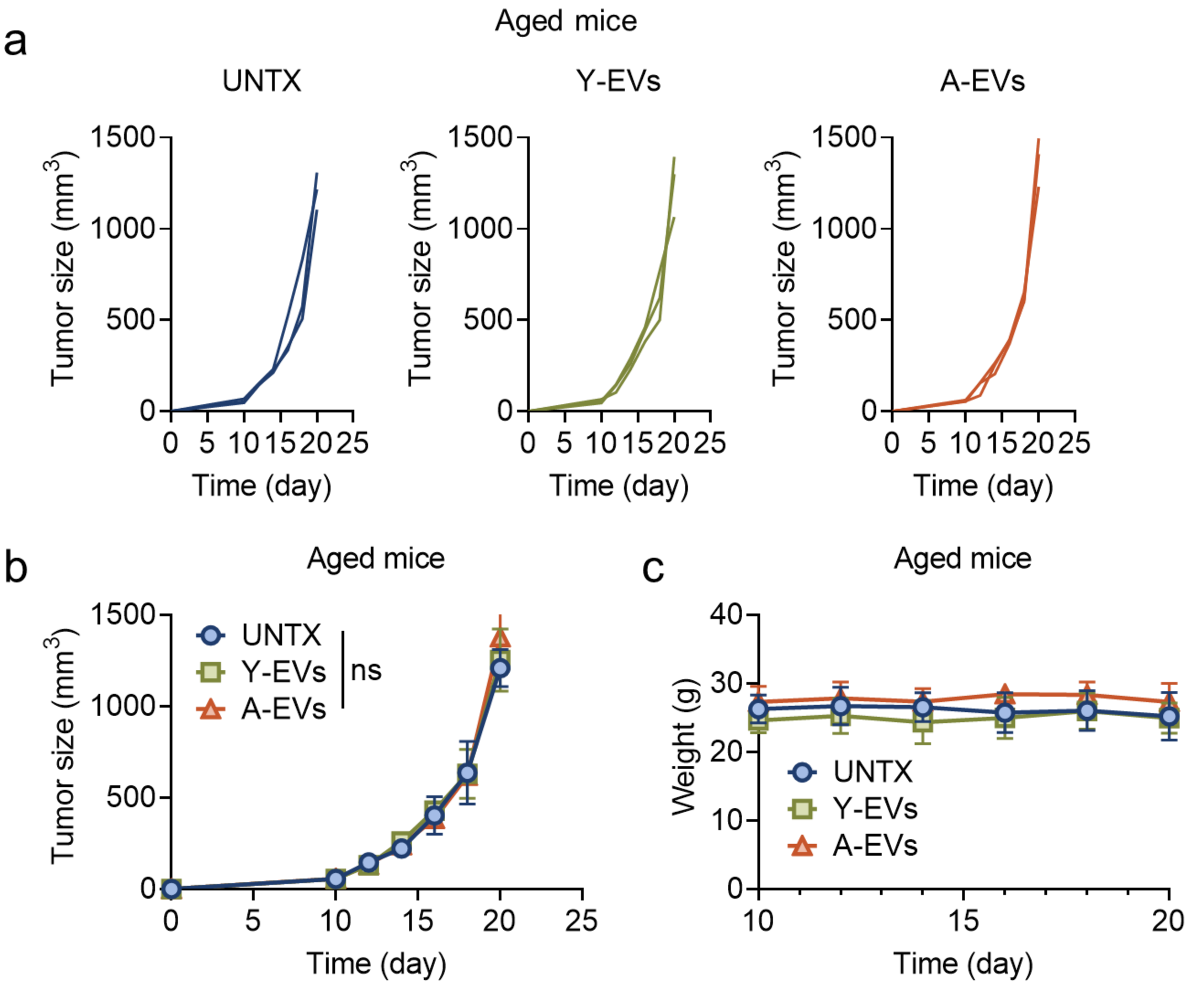


**Figure S4**. Therapeutic efficacy in B16F10 tumor-bearing aged mice (18-month-old) in different groups, including untreated mice, Y-EV-injected mice, and A-EV-injected mice (n=3). a, The tumor growth curves of B16F10 tumor-bearing aged mice in each group (n=3). b, Average curves of B16F10 tumor-bearing aged mice in each group (n=3). c, The weight curves of B16F10 tumor-bearing aged mice (n=3). ns. $P$>0.05. The data are presented as mean ±s.d.

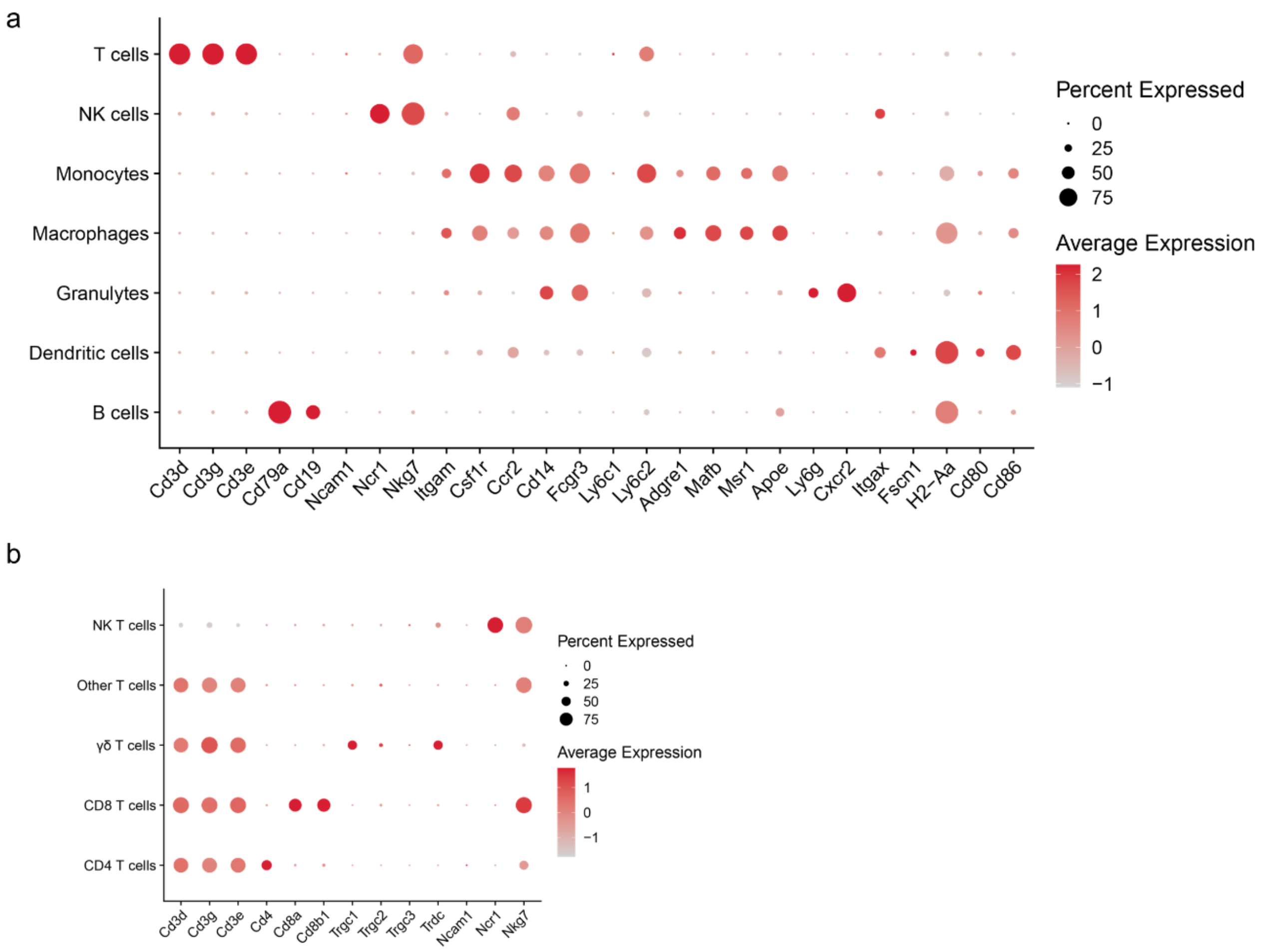


**Figure S5**. Differential expression of genes in scRNA-seq. a, Dot plot of immune cell marker genes. b, Dot plot of T cell marker genes, including *Cd3d*, *Cd3g*, *Cd3e*, *Cd4*, *Cd8a*, *Cd8b1*, *Trgc1*, *Trgc2*, *Trgc3*, *Trdc*, *Ncam1*, *Ncr1*, *Nkg7*.

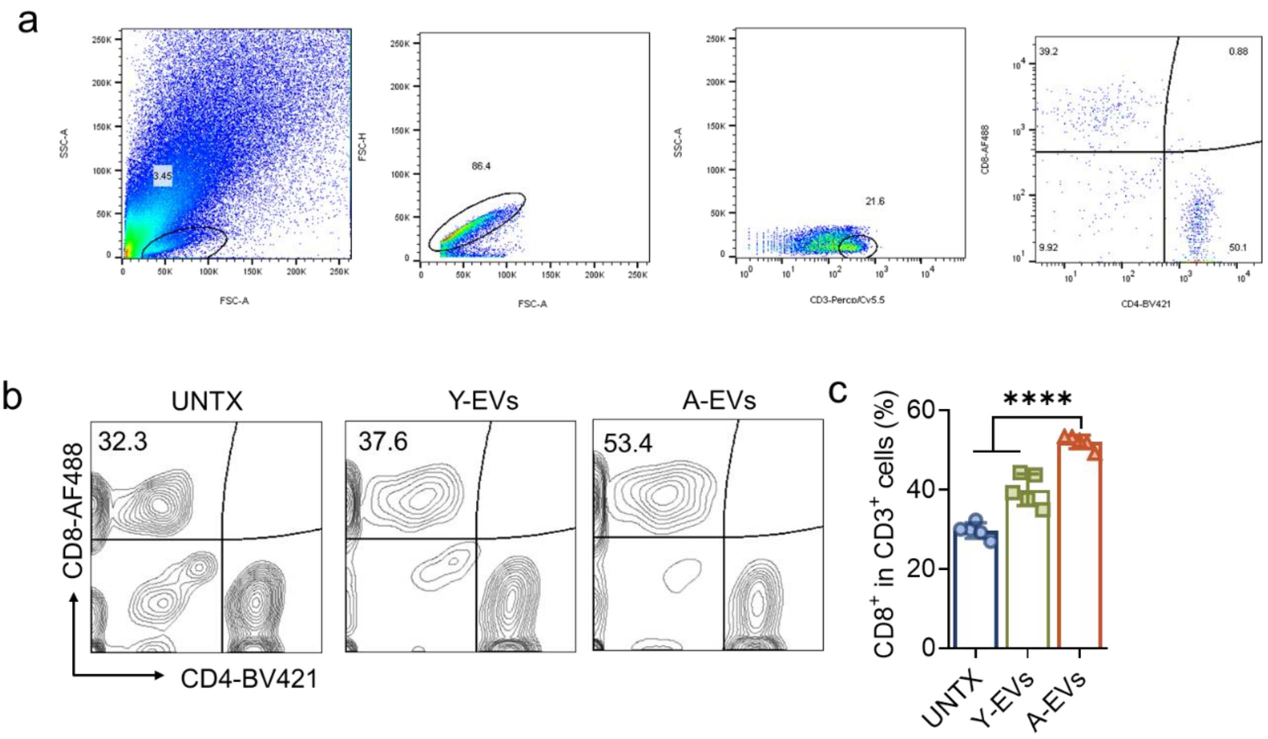


**Figure S6**. Flow cytometry of T cells in the tumor site. a, flow cytometry gating strategy of T cells in the tumor site. b, Representative flow cytometry plots of $CD8^+$ T cells in $CD3^+$ T cells of mice from UNTX, Y-EVs, and A-EVs groups. c, The corresponding quantitative results of $CD8^+$ T cells in (b) (n=5). Statistical significance was determined by one-way ANOVA with the Tukey post hoc test. ****$P$<0.0001, ***$P$ <0.001. The data are presented as mean ± s.d.

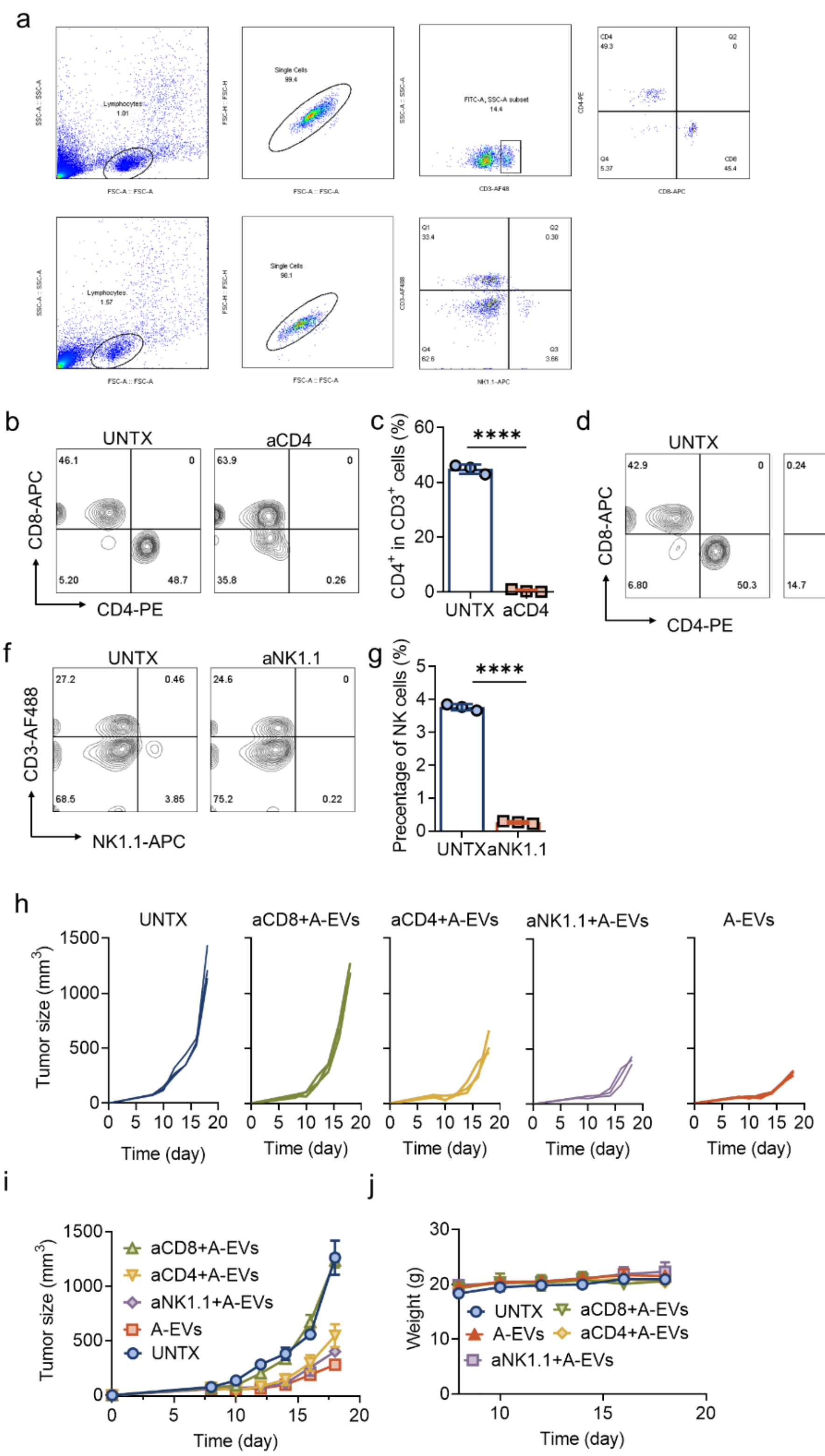


**Figure S7**. Therapeutic efficacy of A-EVs after neutralizing the corresponding antibodies *in vivo*. a, Flow cytometry gating strategy of T cells, NK cells in PBMCs. b, Representative flow cytometry plots of $CD4^+$ T cells in $CD3^+$ T cells of mice from UNTX and aCD4 group. c, The corresponding quantitative results of $CD4^+$ T cells in (b) (n=3). d, Representative flow cytometry plots of $CD8^+$ T cells in $CD3^+$ T cells of mice from UNTX and aCD8 group. e, The corresponding quantitative results of $CD8^+$ T cells in (d) (n=3). f, Representative flow cytometry plots of $NK1.1^+$ cells of mice from UNTX and aNK1.1 group. g, The corresponding quantitative results of NK cells in (f) (n=3). h, Tumor growth curves of B16F10 tumor-bearing mice in each group, including UNTX, aCD4+A-EVs, aCD8+A-EVs, aNK1.1+A-EVs and A-EVs (n=3). i, The average curves of B16F10 tumors in each group (n=3). k, Weight of tumor-bearing mice during therapy (n=3). Statistical significance was determined by one-way ANOVA with the Tukey post hoc test. ****$P$<0.0001. The data are presented as mean ± s.d.

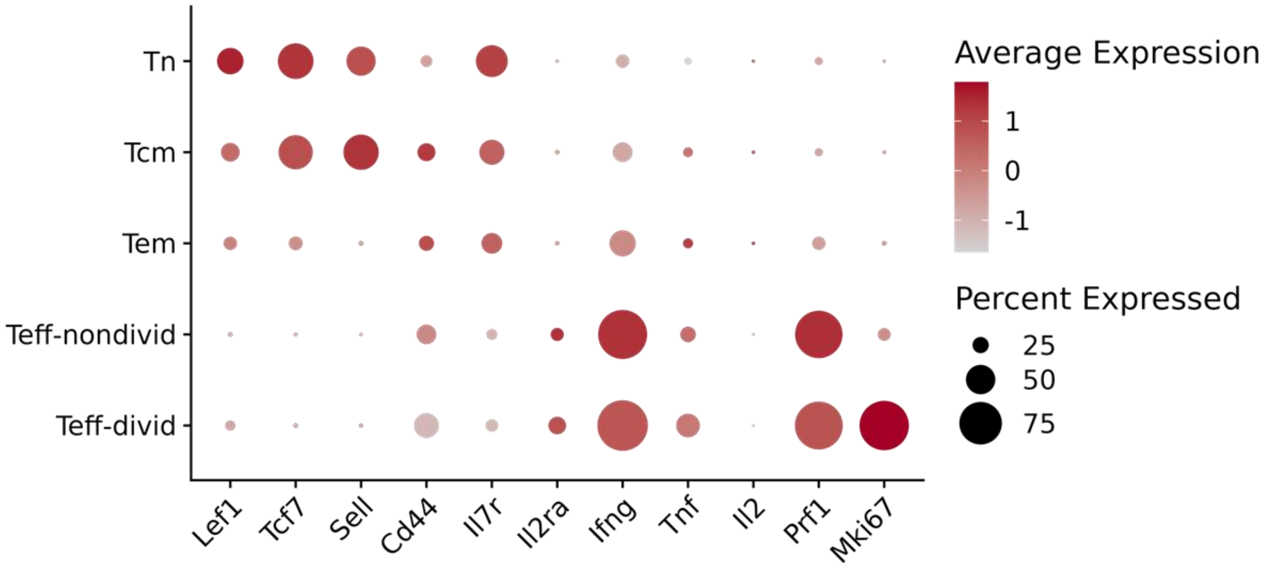


**Figure S8**. Dot plot of $CD8^+$ marker gene expression.

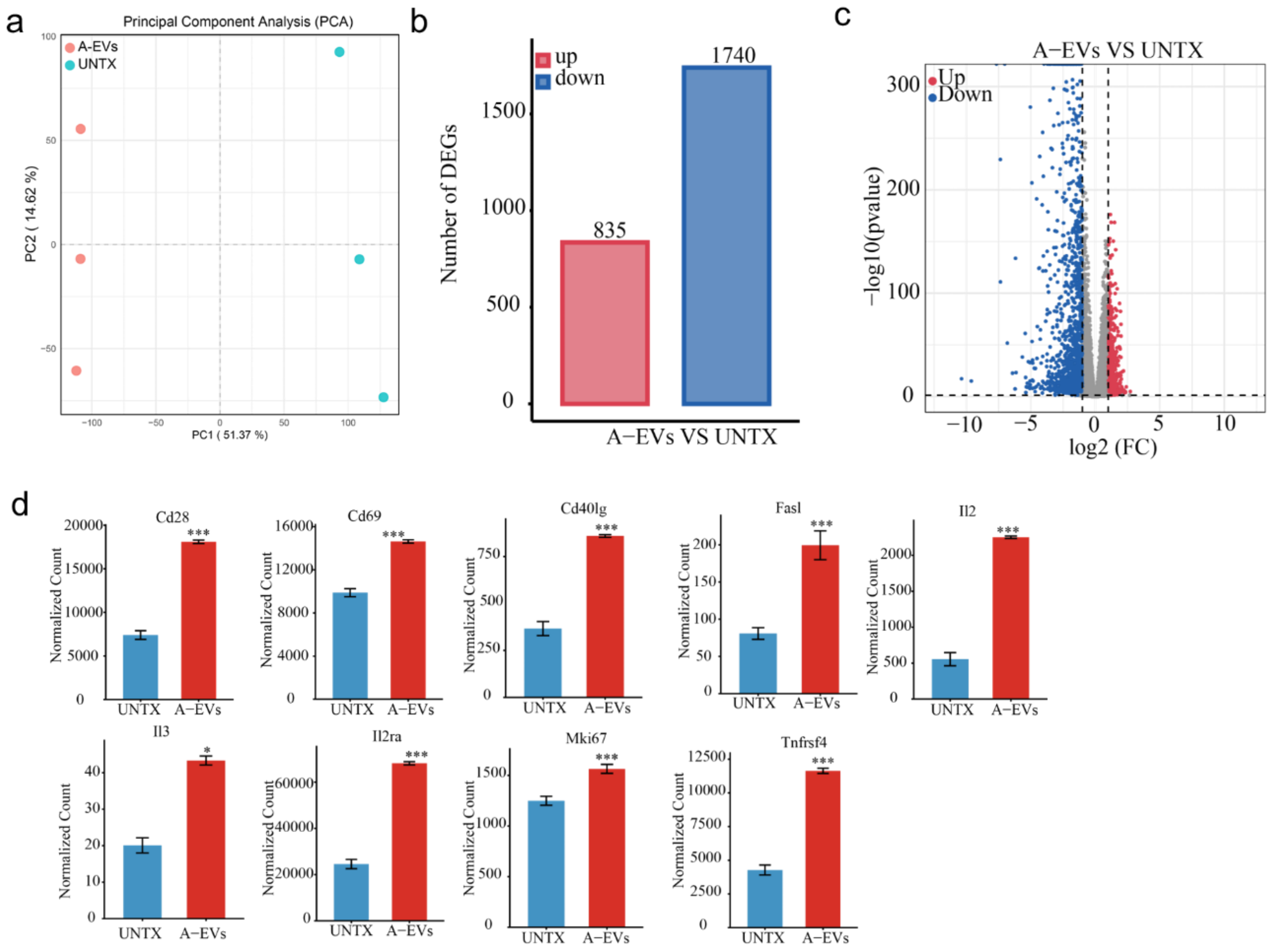


**Figure S9**. RNA-seq of $CD8^+$ $CD44^+$ T cells cocultured with A-EVs *in vitro*. a, PCA analysis of A-EV-treated $CD8^+$ $CD44^+$ T cells and untreated $CD8^+$ $CD44^+$ T cells (n=3). b, Bar plot of differential genes between A-EV-treated $CD8^+$ $CD44^+$ T cells and untreated $CD8^+$ $CD44^+$ T cells. c, Volcano plot of differential expression genes between A-EV-treated $CD8^+$ $CD44^+$ T cells and untreated $CD8^+$ $CD44^+$ T cells. d, Bar plots of differential expression genes between A-EV-treated $CD8^+$ $CD44^+$ T cells and untreated $CD8^+$ $CD44^+$ T cells, including *Il2Ra, Cd40lg, Fasl, Il2, Il3, Cd28, Cd69, Mki67, Tnfrsf4* (n=3). Statistical significance was determined by Student's t-test (two-tailed). ***$P$ <0.001, *$P$<0.05. The data are presented as mean ±s.d.

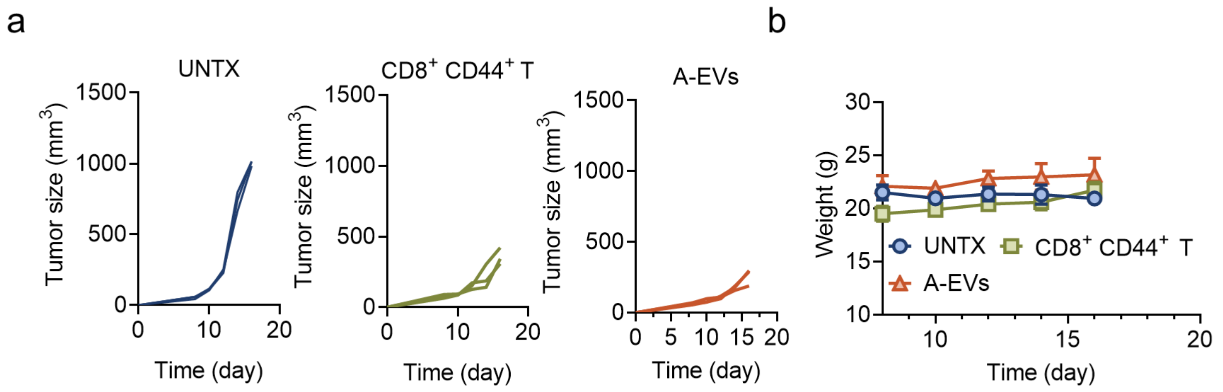


**Figure S10**. The reinfusion of CD8$^+$ CD44$^+$ T cells delays the tumor growth. a, The curves of tumor volume in each group (n=3). b, The weight curve of tumor-bearing mice in each group (n=3). The data are presented as mean ±s.d.

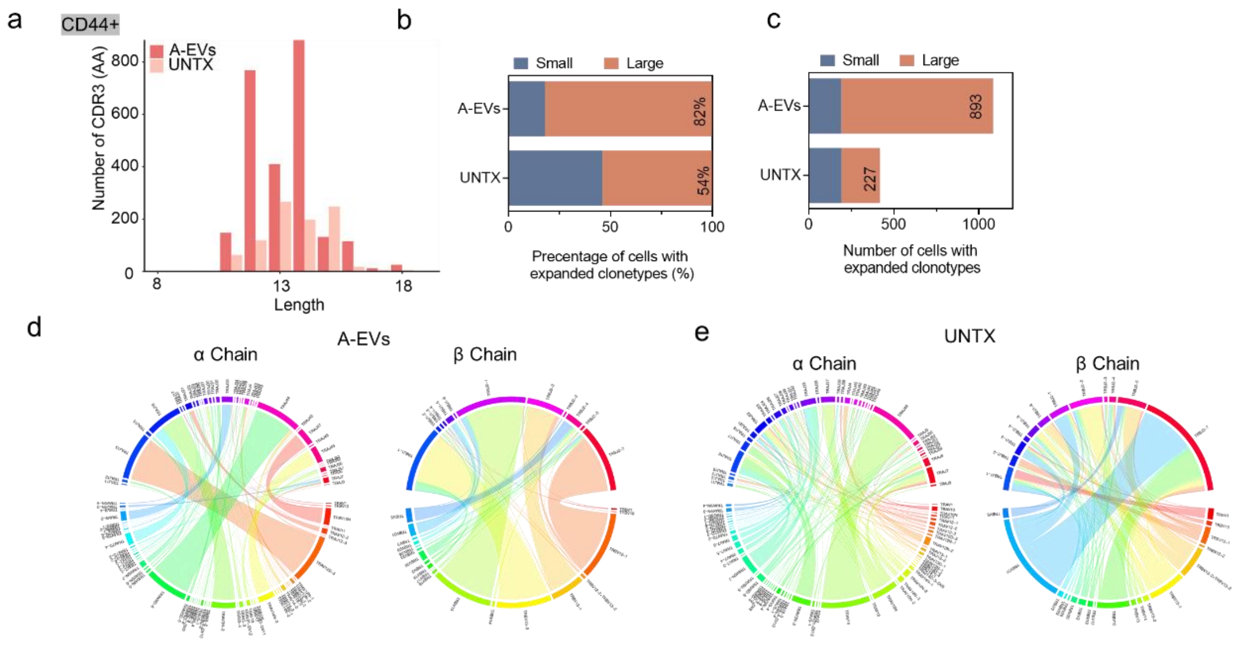


**Figure S11**. TCR analysis of CD8$^+$ CD44$^+$ T cells from A-EV-treated tumor and untreated tumor. a, Bar plots of the number distribution of CDR3 in CD8$^+$ CD44$^+$ T cells in different groups. b, The percentage of different clonotypes of CD8$^+$ CD44$^+$ T cells (Small: clonotype <10, Large: clonotype ≥10). c, Absolute number of different clonotypes of CD8$^+$ CD44$^+$ T cells. d, Circle plots of TCR sequence of CD8$^+$ CD44$^+$ T cells from A-EV-treated tumor (left: α chain, right: β chain). e, Circle plots of TCR sequence of CD8$^+$ CD44$^+$ T cells from untreated tumor (left: α chain, right: β chain).

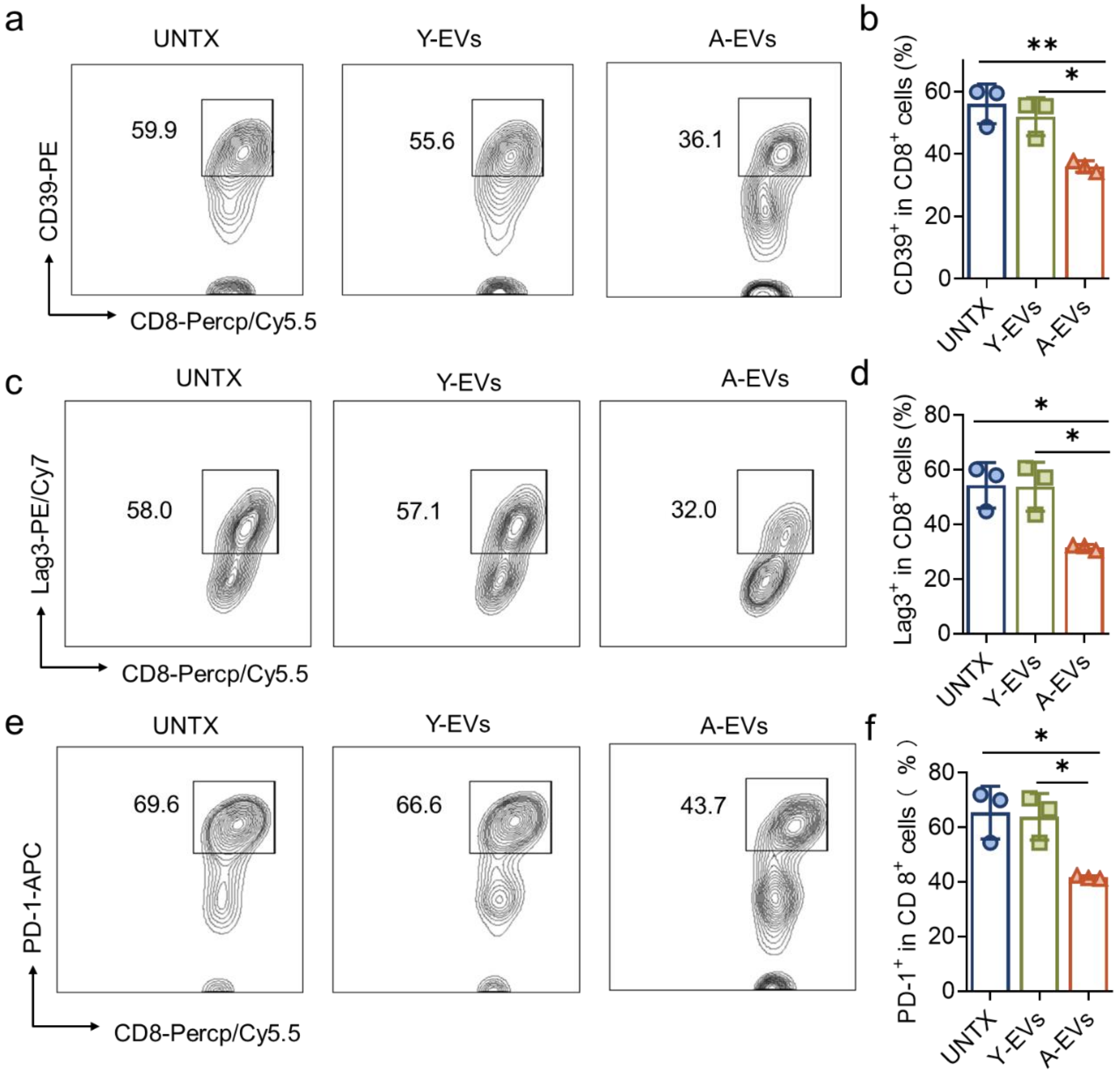


**Figure S12**. Bystander-related markers in the tumor site. a, Representative flow cytometry plots of the expression of CD39 on CD8$^+$ T cells in tumors of mice from each group. b, Corresponding quantitative analysis of (a) (n=3). c, Representative flow cytometry plots of the expression of Lag3 on CD8$^+$ T cells in tumors of mice from each group. d, Corresponding quantitative analysis of (c) (n=3). e, Representative flow cytometry plots of the expression of PD-1 on CD8$^+$ T cells in tumors of mice from each group. f, Corresponding quantitative analysis of (e) (n=3). Statistical significance was determined by one-way ANOVA with the Tukey post hoc test. ***P*<0.01, * *P*<0.05. The data are presented as mean ± s.d.

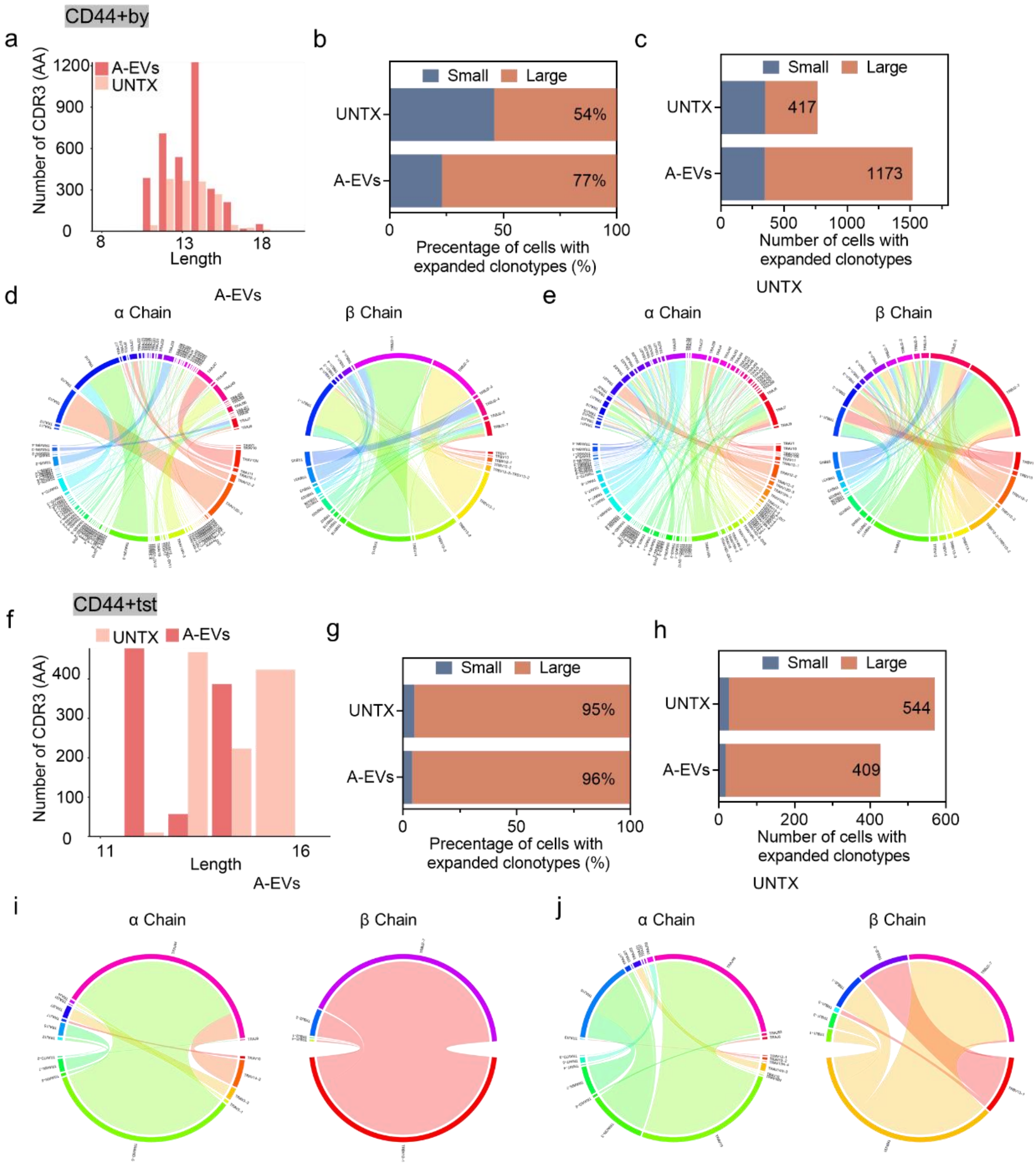


**Figure S13**. TCR analysis of bystander T cells and tumor-specific T cells from CD8$^+$ CD44$^+$ T cells. a, Bar plots of the number distribution of CDR3 in CD8$^+$ CD44$^+$ T bystander cells in different groups. b, Percentage of different clonotypes in CD8$^+$ CD44$^+$ T bystander cells (Small: clonotype <10, Large: clonotype ≥10). c, Absolute number of different clonotypes in CD8$^+$ CD44$^+$ T bystander cells. d, Circle plots of TCR sequence of CD8$^+$ CD44$^+$ T bystander cells from A-EV-treated tumor (left: α chain, right: β chain). e, Circle plots of TCR sequence of CD8$^+$ CD44$^+$ T bystander cells from untreated tumor (left: α chain, right: β chain). f, Bar plots of the number distribution of CDR3 in

CD8$^+$ CD44$^+$ tumor-specific T cells in different groups. g, Percentage of different clonotypes of CD8$^+$ CD44$^+$ tumor-specific T cells (Small: clonotype <10, Large: clonotype ≥10). h, Absolute number of different clonotypes of CD8$^+$ CD44$^+$ tumor-specific T cells. i, Circle plots of TCR sequence of CD8$^+$ CD44$^+$ tumor-specific T cells from A-EV-treated tumor (left: α chain, right: β chain). j, Circle plots of TCR sequence of CD8$^+$ CD44$^+$ tumor-specific T cells from untreated tumor (left: α chain, right: β chain).

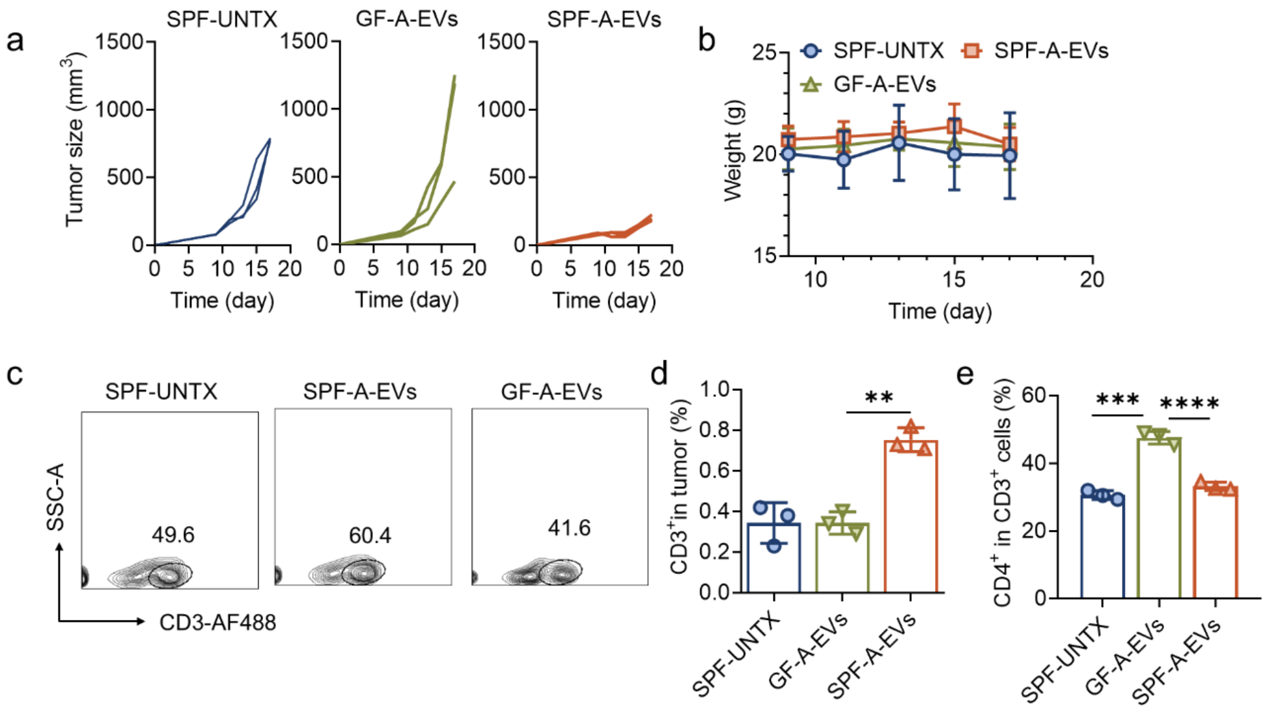


**Figure S14.** Therapeutic efficacy of A-EVs on mice living in different cleanliness levels. a, Curves of tumor volume of each group, including SPF-UNTX, SPF-A-EVs, and GF-A-EVs (n=3). b, The weight curve of tumor-bearing mice in each group (n=3). c, Representative flow cytometry plots of $CD3^+$ T cells in tumors from mice in SPF-UNTX, SPF-A-EVs, and GF-A-EVs groups. d, Corresponding quantitative results of (c) (n=3). e, Corresponding quantitative results of Fig. 3n (n=3). Statistical significance was determined by Student's t-test (two-tailed) and one-way ANOVA with the Tukey post hoc test. ****$P<0.0001$, **$P<0.01$. The data are presented as mean ± s.d.

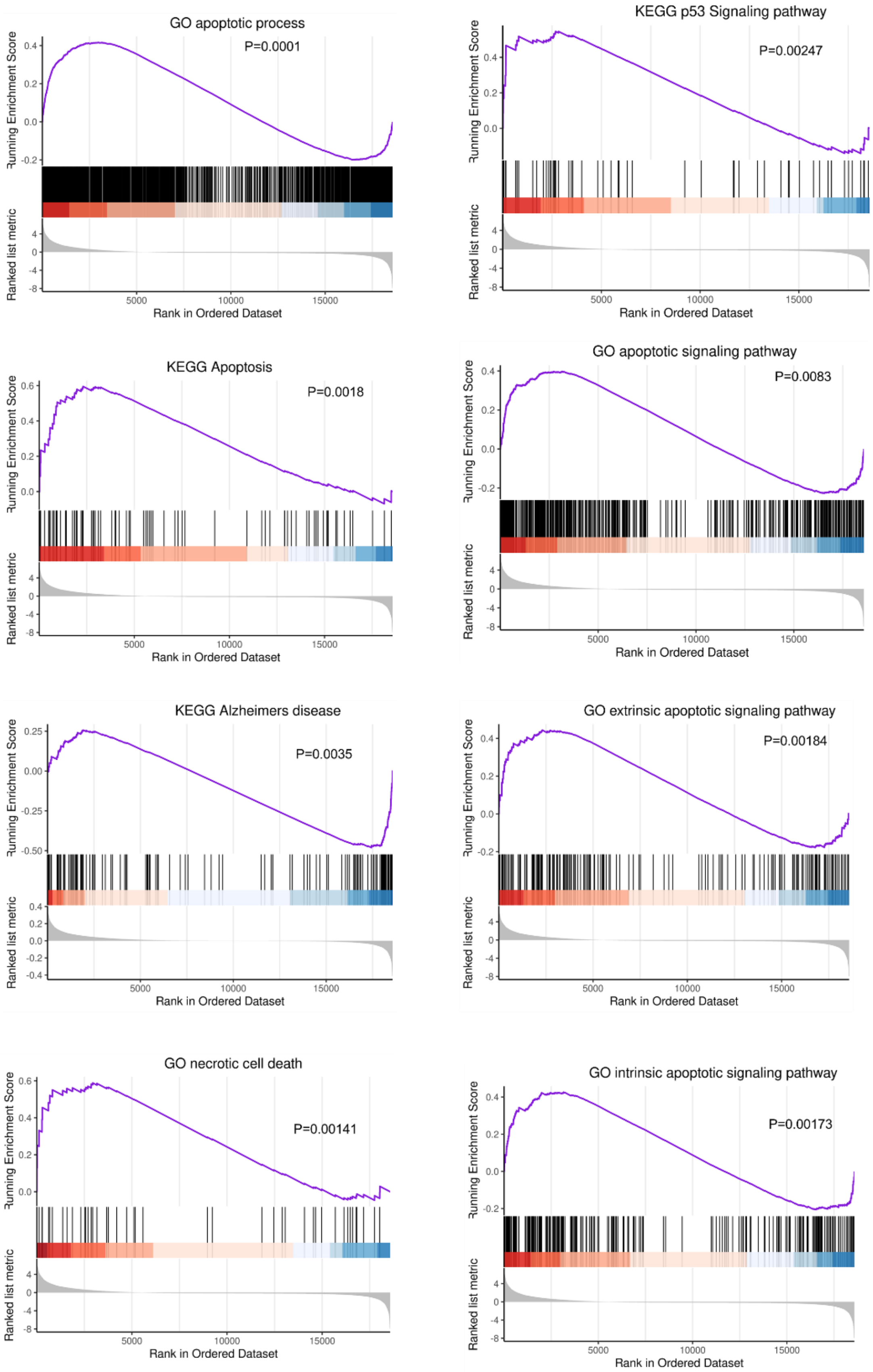


**Figure S15**. KEGG and GO pathway enrichment analysis of comparative analysis between malignant cells from A-EV-treated tumor and malignant cells from untreated tumor.

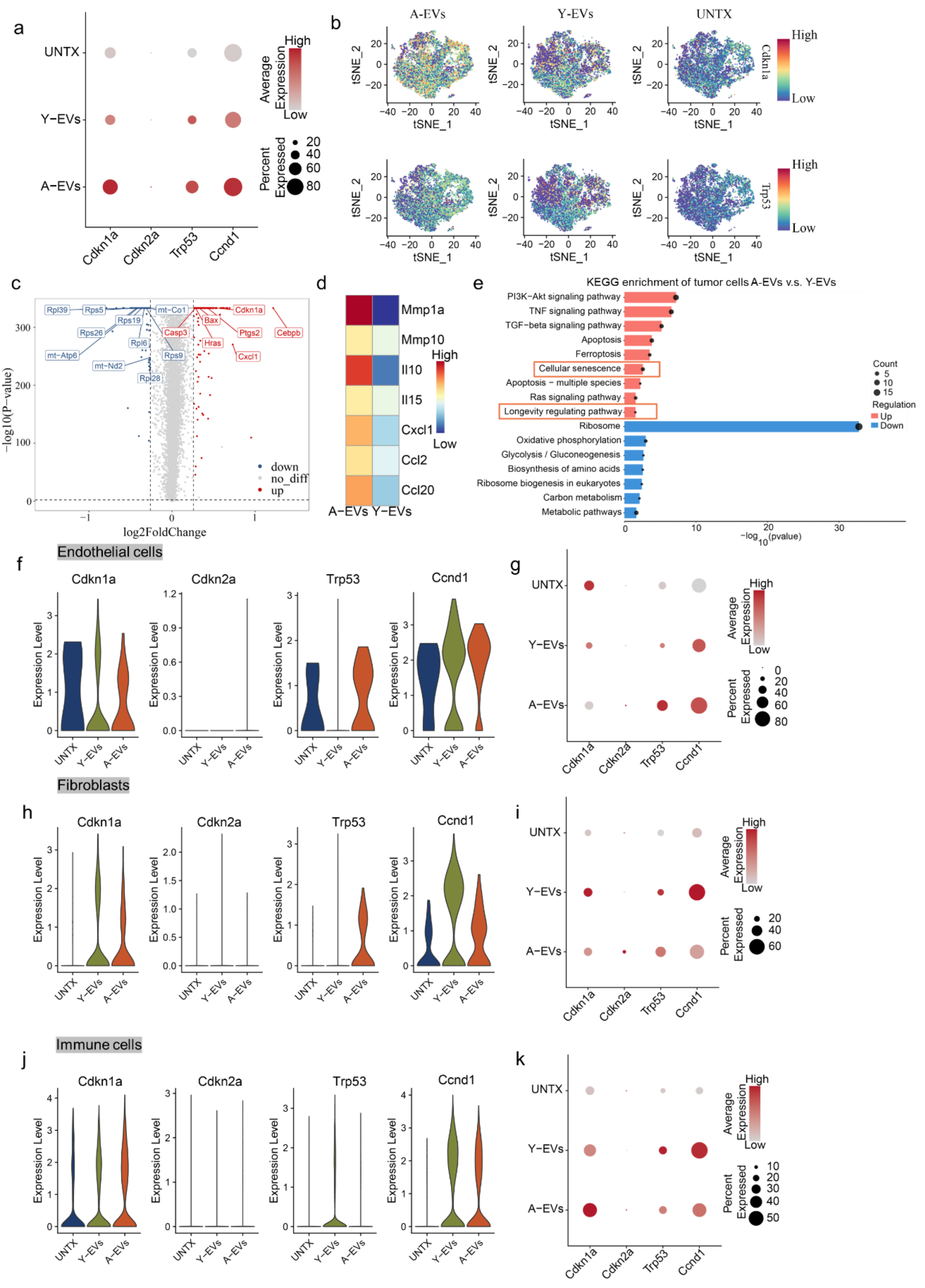
a
UNTX
Y−EVs
A−EVs
Cdkn1a
Cdkn2a
Trp53
Ccnd1
Average Expression
High
Low
Percent Expressed
20
40
60
80
b
A-EVs
Y-EVs
UNTX
tSNE_1
tSNE_2
Cdkn1a
Trp53
High
Low
c
Rpl39
Rps5
mt−Co1
Cdkn1a
Rps19
Bax
Rps26
Casp3
Ptgs2
Cebpb
Rpl6
Hras
mt−Atp6
Rps9
Cxcl1
mt−Nd2
Rpl28
down
no_diff
up
log2FoldChange
−log10(P−value)
d
Mmp1a
Mmp10
Il10
Il15
Cxcl1
Ccl2
Ccl20
High
Low
A−EVs Y−EVs
e
KEGG enrichment of tumor cells A-EVs v.s. Y-EVs
PI3K−Akt signaling pathway
TNF signaling pathway
TGF−beta signaling pathway
Apoptosis
Ferroptosis
Cellular senescence
Apoptosis − multiple species
Ras signaling pathway
Longevity regulating pathway
Ribosome
Oxidative phosphorylation
Glycolysis / Gluconeogenesis
Biosynthesis of amino acids
Ribosome biogenesis in eukaryotes
Carbon metabolism
Metabolic pathways
Count
5
10
15
Regulation
Up
Down
−log10(pvalue)
Endothelial cells
f
g
Expression Level
Fibroblasts
h
i
Immune cells
j
k

**Figure S16**. scRNA-seq analysis of tumors after the treatment of A-EVs and Y-EVs. a, Dot plot of different expression of genes in malignant cells from UNTX, Y-EV, and A-EV-treated tumor. b, T-SNE mapping of the expression of *Cdkn1a* and *trp53* in malignant cells from UNTX, Y-EV, and A-EV-treated tumor. c, Volcano plots of differential expression genes compared between malignant cells from A-EV-treated tumor and tumor cells from Y-EV-treated tumor. d, Heatmap of differential expression genes related to SASPs of malignant cells. e, KEGG enrichment bar plot of malignant cells, compared between malignant cells from A-EV-treated tumor and malignant cells from Y-EV-treated tumor. f, Violin plot of differential expression of genes related to senescence in endothelial cells from UNTX, Y-EV, and A-EV-treated tumor. g, Dot plot of differential expression of genes related to senescence in endothelial cells from UNTX, Y-EV, and A-EV-treated tumor. h, Violin plot of differential expression of genes related to senescence in endothelial cells from UNTX, Y-EV, and A-EV-treated tumor. i, Dot plot of differential expression of genes related to senescence in fibroblasts from UNTX, Y-EV, and A-EV-treated tumor. j, Violin plot of differential expression of genes related to senescence in immune cells from UNTX, Y-EV, and A-EV-treated tumor. k, Dot plot of differential expression of genes related to senescence in immune cells from UNTX, Y-EV, and A-EV-treated tumor.

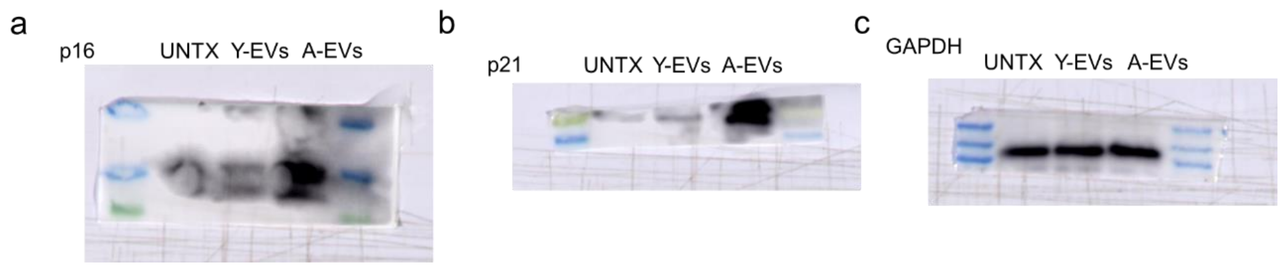


**Figure S17**. Western blot analysis of p21, p16, and GAPDH in tumors from mice in UNTX, Y-EVs, and A-EVs groups.

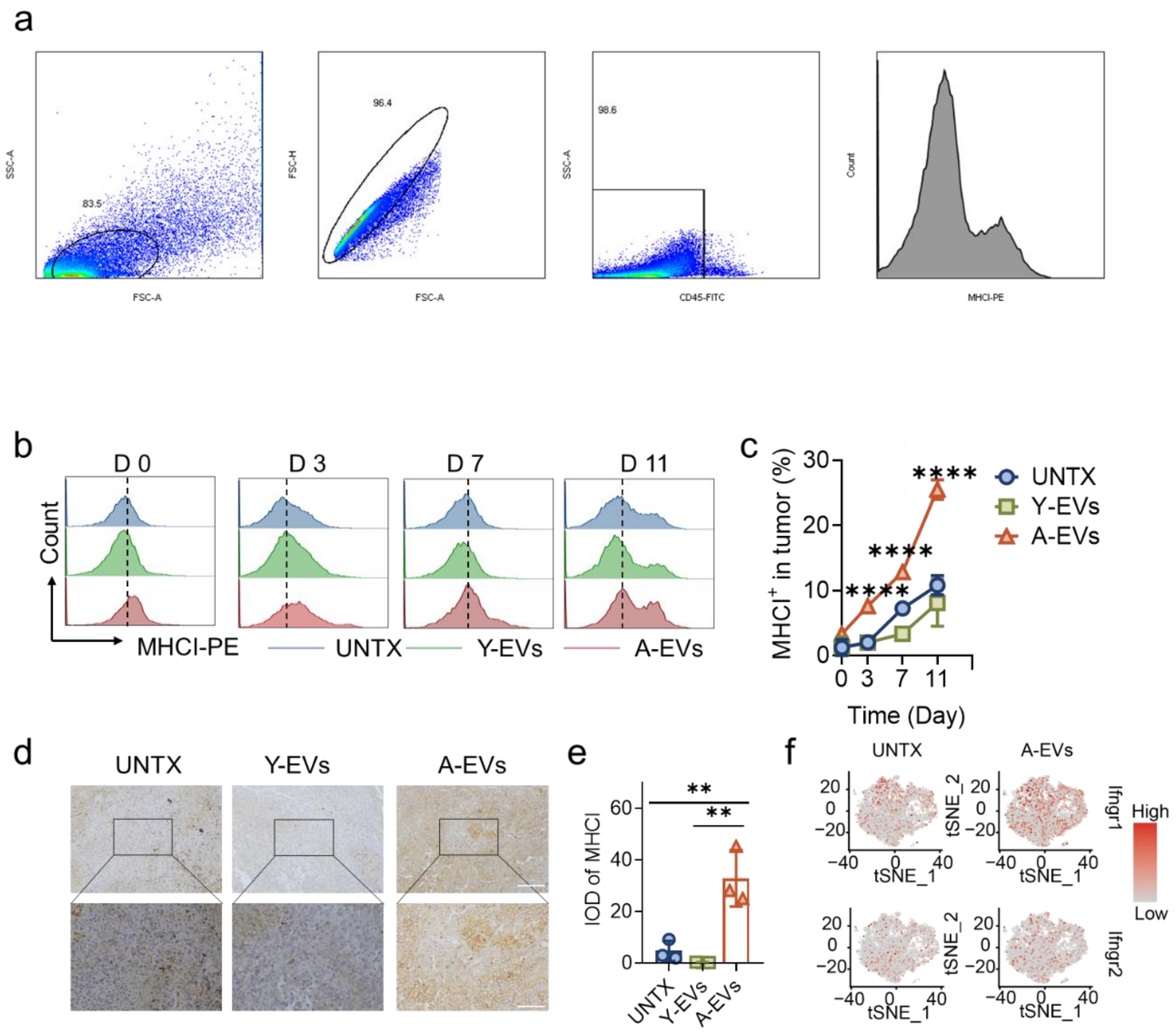


**Figure S18**. The enhanced expression of MHCI in tumor cells after the treatment of A-EVs. a, Flow cytometry gating strategy of analyzing tumor cells in various treated tumors. b, Representative flow cytometry plots of MHC-I$^{+}$ in tumor cells of mice from UNTX, Y-EVs, and A-EVs groups. c, The corresponding quantitative results of (b) over time (n=3). d, Representative images of MHC-I immunohistochemical staining of tumor sections from mice in UNTX, Y-EVs, and A-EV group (Scale bar: 100 μm (up); 50 μm (down). e, The quantitative results of (d) (n=3). f, Highlighted t-SNE plot of *Ifngr1* and *Ifngr2* of malignant cells. Statistical significance was determined by one-way ANOVA with the Tukey post hoc test. ****$P$<0.0001, **$P$<0.01. The data are presented as mean ± s.d.

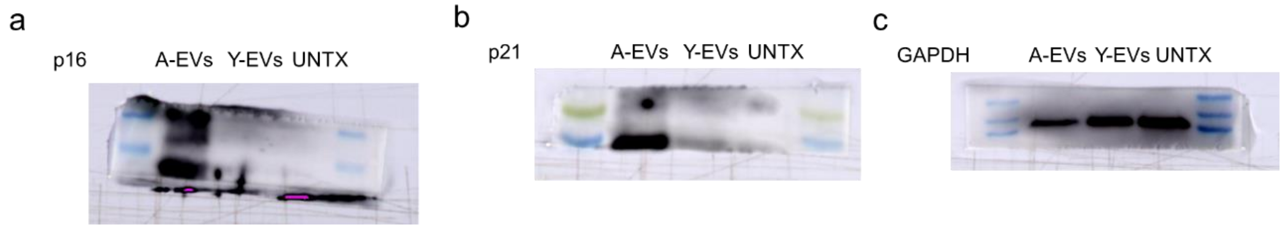


**Figure S19**. Western blot analysis of p21, p16, and GAPDH in tumor cells cocultured with A-EVs, Y-EVs for 48 h.

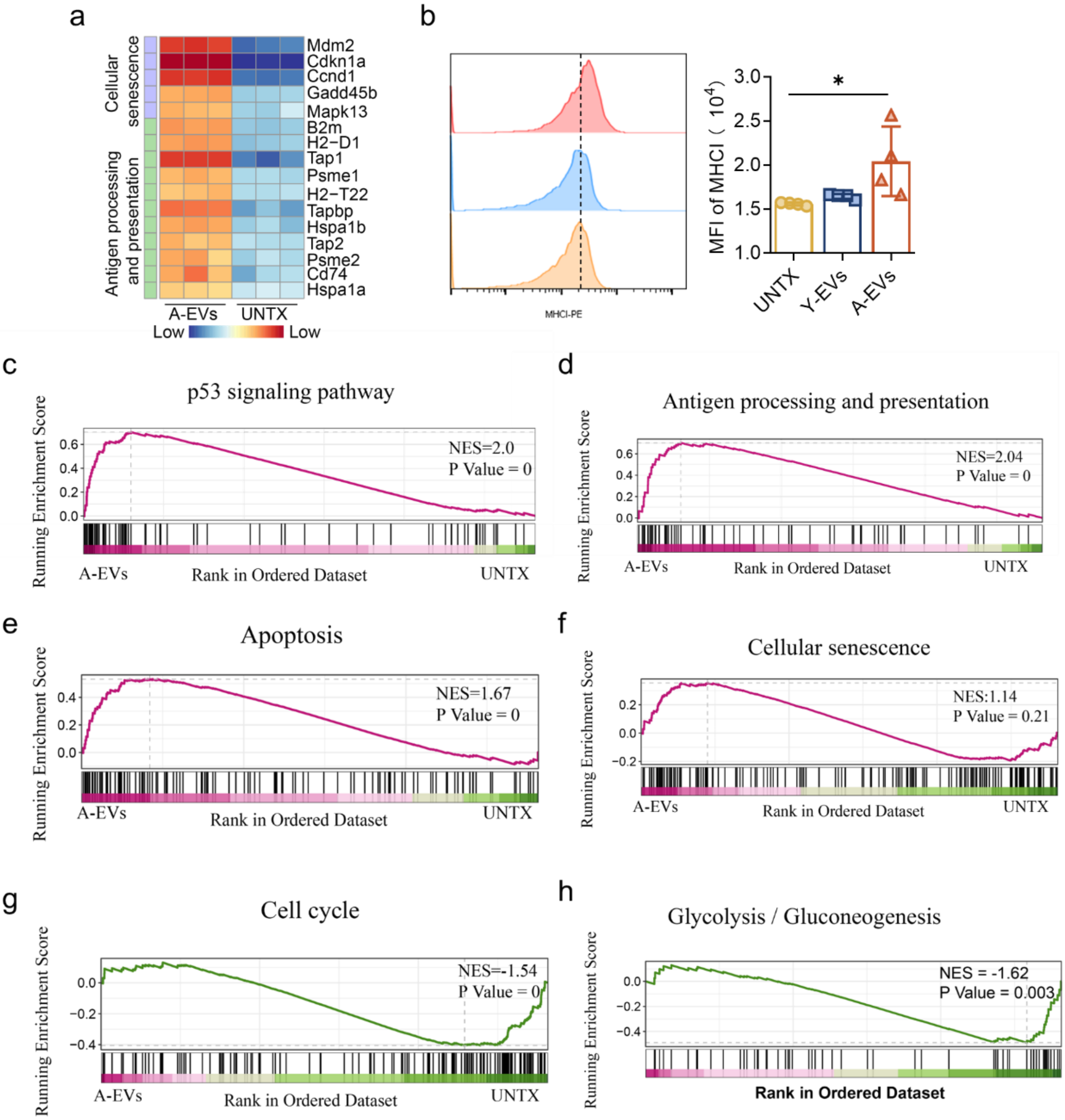


**Figure S20**. RNA-seq of tumor cells after the treatment of A-EVs *in vitro*. a, Heatmap of differential expression of genes related to cellular senescence and antigen processing and presentation in tumor cells after the treatment of A-EVs *in vitro*. b, Representative flow cytometry plots of MHC-I$^{+}$ in tumor cells after the treatment of A-EVs, Y-EVs *in vitro*, and the corresponding quantitative results (n=4). c-h, KEGG, and GO pathway enrichment analysis of tumor cells, compared between A-EV-treated tumor cells and untreated tumor cells. Statistical significance was determined by one-way ANOVA with the Tukey post hoc test. *$P$<0.05. The data are presented as mean ± s.d.

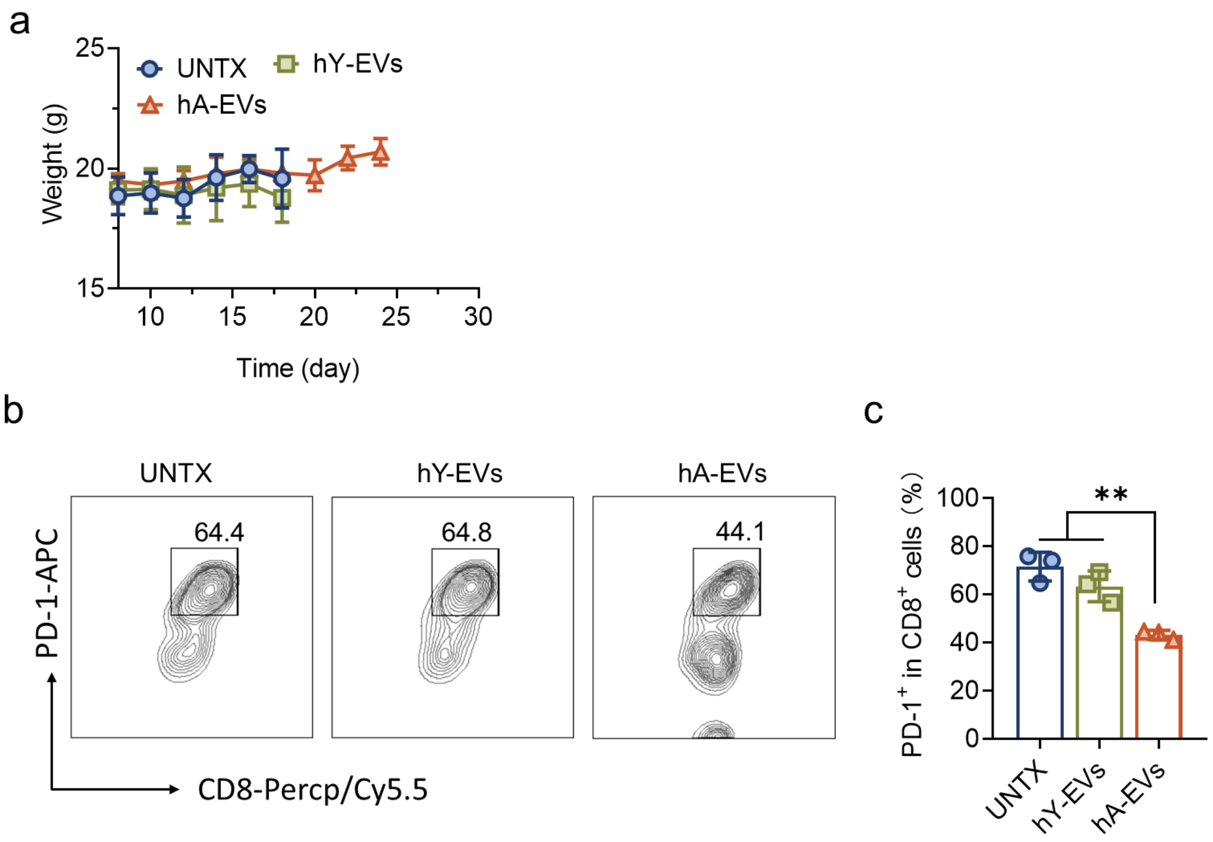


**Figure S21**. Human aged EVs inhibited tumor growth in mice. a, The weight curve of tumor-bearing mice in each group (n=5). b, Representative flow cytometry plots of the expression of PD-1 on $CD8^+$ T cells in tumors of mice from each group. c, Corresponding quantitative analysis of (b) (n=3). Statistical significance was determined by one-way ANOVA with the Tukey post hoc test. ***P*<0.01. The data are presented as mean ±s.d.

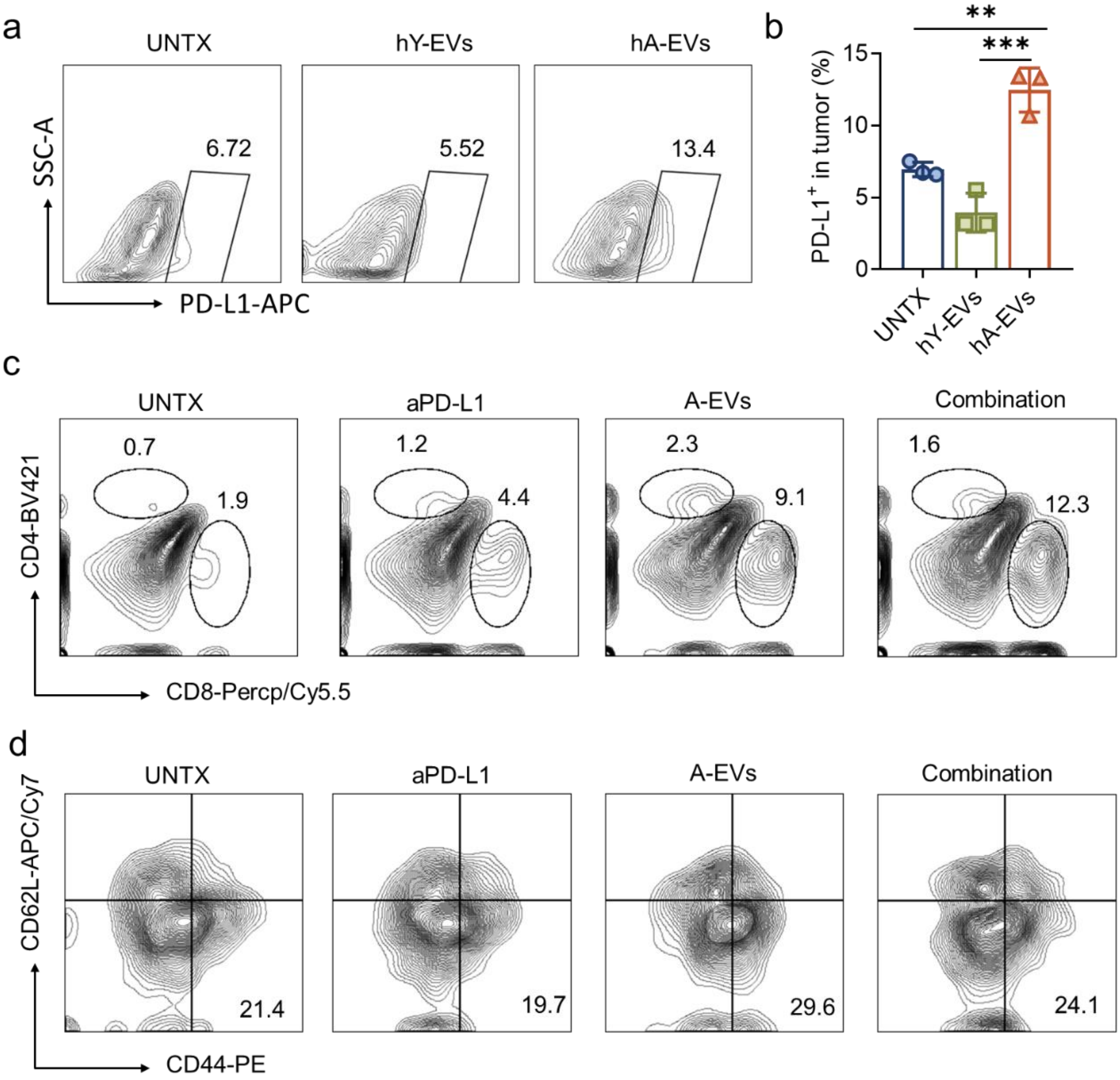


**Figure S22**. Combination therapy of A-EVs and aPD-L1. a, Representative flow cytometry plots of PD-L1 in tumors of mice from each group. b, Corresponding quantitative analysis of PD-L1 level in tumors of mice after the treatment of human EVs (n=3). c, Representative flow cytometry plots of $CD4^+$ and $CD8^+$ T cells in tumors of mice from each group. d, Representative flow cytometry plots of effector memory and effector T cells (Tem/eff) among $CD8^+$ T cells from the tumors in each group. Statistical significance was determined by one-way ANOVA with the Tukey post hoc test. ***$P$<0.001, **$P$<0.01. The data are presented as mean ± s.d.
